# Cell-Penetrating D-Peptides Retain Antisense Morpholino Oligomer Delivery Activity

**DOI:** 10.1101/2021.09.30.462617

**Authors:** Carly K. Schissel, Charlotte E. Farquhar, Annika B. Malmberg, Andrei Loas, Bradley L. Pentelute

## Abstract

Cell-penetrating peptides (CPPs) can cross the cell membrane to enter the cytosol and deliver otherwise non-penetrant macromolecules such as proteins and oligonucleotides. For example, recent clinical trials have shown that a CPP attached to phosphorodiamidate morpholino oligomers (PMO) resulted in higher muscle concentration, increased exon-skipping and dystrophin production relative to another study of the PMO alone in patients of Duchenne muscular dystrophy. Therefore, effective design and study of CPPs could help enhance therapies for difficult-to-treat diseases. So far, the study of CPPs for PMO delivery has been restricted to predominantly canonical L-peptides. We hypothesized that mirror-image D-peptides could have similar PMO delivery activity as well as enhanced proteolytic stability, facilitating their characterization and quantification from biological milieu. We found that several enantiomeric peptide sequences could deliver a PMO-biotin cargo with similar activities, while remaining stable against serum proteolysis. The biotin label allowed for affinity capture of fully intact PMO-peptide conjugates from whole cell and cytosolic lysates. By profiling a mixture of these constructs in cells, we determined their relative intracellular concentrations. When combined with PMO activity, these concentrations provide a new metric for delivery efficiency which may be useful for determining which peptide sequence to pursue in further pre-clinical studies.

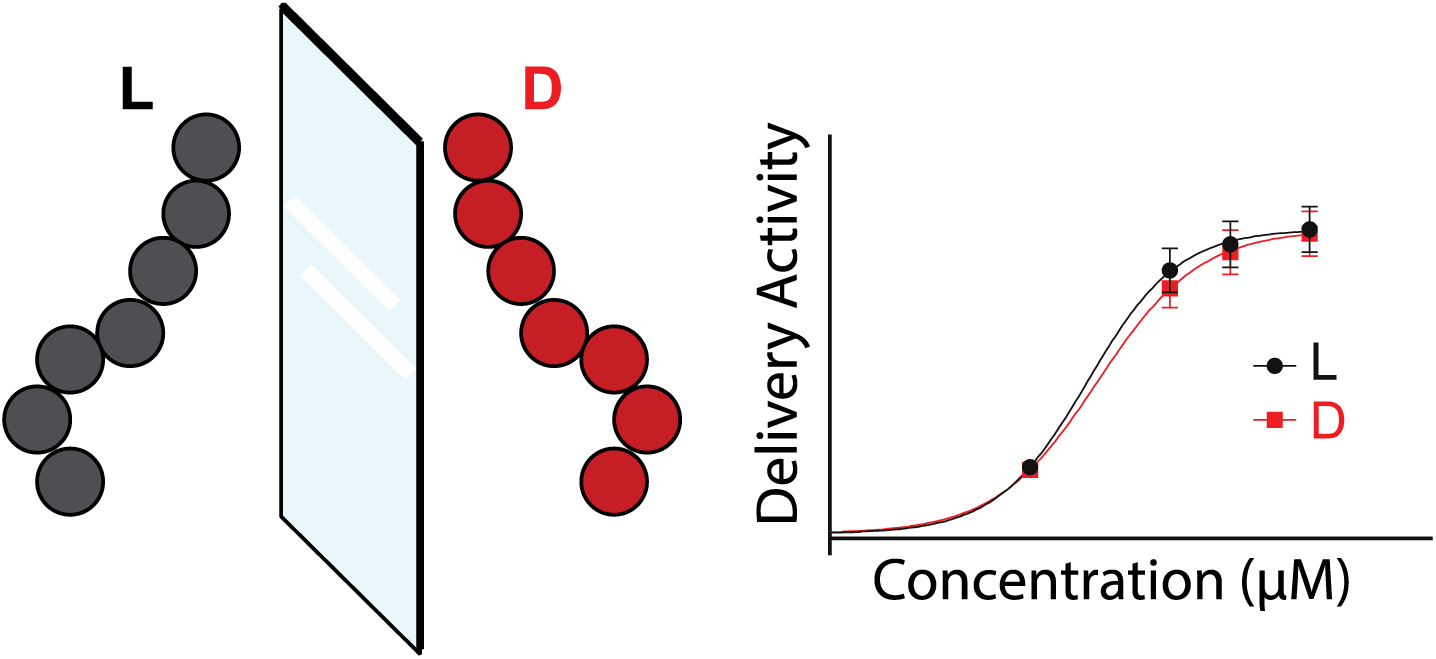

## Introduction

Cell-penetrating peptides (CPPs) can help treat disease by enhancing the delivery of cell-impermeable cargo. CPPs are a class of peptides that are capable of directly entering the cell cytosol.^1–3^ These sequences can deliver covalently bound cargo, offering therapeutic potential to macromolecules otherwise restricted to extracellular targets. Although CPPs have been widely studied since their discovery, the field lacks robust methodology to quantify cell entry and penetration efficacy. This dearth of knowledge is due to the complicated mechanisms of CPP cell entry and the many variables that affect CPP efficacy in any given assay—such as peptide concentration, cell type, temperature, treatment time, and cargo.^4^ For example, for the well-studied CPP penetratin (RQIKIWFQNRRMKWKK), the reported ratio between intracellular and extracellular concentration ranges from 0.6:1.0 to 95.0:1.0.^5,6^ In addition, it is challenging to determine subcellular localization once a peptide is internalized, despite advances in fluorescence, immunoblot, and mass spectrometry detection.^7^ The choice of CPP cargo adds an additional confounding factor, as studies in our laboratory have demonstrated that the cell-penetrating ability of more than ten common CPPs differs when bound to a cyanine dye versus a macromolecular drug, with no discernable trend.^8^ Therefore, effective development of CPPs requires new methodology for understanding CPP cell entry and subcellular localization that can be carried out on the CPP-cargo conjugate.

A therapeutic macromolecule that would benefit from enhanced delivery is phosphorodiamidate morpholino oligomer (PMO), which has recently reached the market as an antisense “exon skipping” therapy for Duchenne muscular dystrophy (DMD). One of the drugs, eteplirsen, is a 10 kDa synthetic antisense oligomer that must reach the nucleus and bind pre-mRNA for its therapeutic effect. However, studies have shown that two-thirds of eteplirsen is cleared renally within 24 h of administration.^9,10^ Several CPPs have been shown to increase PMO uptake, and recent clinical trial results have shown that once-monthly dosing of SRP-5051 resulted in higher muscle concentration, increased exon-skipping and dystrophin production at 12 weeks as compared to once weekly dosing of eteplirsen after 24 weeks in a different study.^11^ Although a wide variety of CPPs have been tested for PMO delivery, they have been limited to the native L-form and studied predominantly with an activity-based assay, forgoing quantitative information on the amount of material inside the cell.^12^

Because PMO is proteolytically stable and invisible to the immune system^10^, attachment to an L-peptide may disrupt the desirable characteristics of PMO; however, conjugates using mirror image peptides may retain these characteristics.^13–15^ D-peptides have been explored as CPPs and have been found to display at times greater activities to their L-counterparts, despite suggested chiral binding interactions between CPPs and the cell membrane.^16,17^ A study on TAT endosomalytic peptide analogs found that the full D-form decreased uptake, but enhanced endosomal escape and proteolytic stability compared to the native form.^15,18^ Another reported that a cyclic D-peptide, when co-administered with insulin, enhanced its oral bioavailability and therapeutic effect in the gut.^19^ Some reports are contentious as to whether mutations to D-amino acids are detrimental to CPP activity, and certainly CPPs that depend on a chiral interaction or highly ordered structure to enter the membrane would lose efficacy from D-amino acid substitutions.^20,21^ Previous studies have suggested that efficient CPPs for PMO delivery lack secondary structure and can enter the cell through clathrin-mediated endocytosis.^22,23^ While it has been found that PMO-D-CPPs have enhanced proteolytic stability over their L-counterparts^24^, their PMO delivery activity has not been investigated. In addition to their underexplored potential, we are interested in studying D-CPPs as potential therapeutic carrier moieties because their fully noncanonical sequence is resistant to proteolysis and may go unrecognized by the host immune system.^15,20^ Despite this potential, mirror image peptides have not yet been fully explored for the delivery of PMO.

The proteolytic stability of mirror image peptides would simplify their characterization after uptake into cells, providing orthogonal information to activity-based assays. Currently, the main method used to characterize PMO-CPP internalization is an in vitro assay in which successful delivery of the active oligomer to the nucleus results in green fluorescence.^25^ While this is an excellent assay to measure PMO-CPP activity, this assay does not give information on the quantity of material inside the cell. Especially for conjugates with a known endocytic mechanism, understanding endosomal escape is crucial. High concentrations of peptides trapped in the endosome would not be apparent by the activity assay alone, but this loss of active peptide could be of great therapeutic detriment.^26^ Therefore, an additional assay that reveals relative quantities of material in different parts of the cell would provide a valuable new metric of CPP delivery, in parallel with our current activity assay. This could be achieved by extracting the cytosolic fraction of treated cells using a mild detergent, such as digitonin, and comparison to the whole cell fraction.^27^ Comparing activity and relative quantity would be a valuable metric for efficiency, where a high efficiency peptide is one with a high ratio of antisense activity to internal concentration.

Existing methods to quantify uptake into cells include fluorescence, immunoblot, and mass spectrometry, but it is still challenging to distinguish between endosomal and cytosolic localization. Several assays can differentiate between endosomal and cytosolic localization using indirect quantification via a readout generated by a delivered cargo, including the chloroalkane penetration assay (CAPA),^28^ GFP complementation assays,^29^ and more recently the NanoClick^30^ assay and SLEEQ^31^ assay. Direct quantification of the cytosolic concentration of a fluorescently-labeled protein construct is possible using fluorescence correlation spectroscopy.^32^

Mass spectrometry is a direct quantification tool that would give information about the concentration of peptides recovered from biological mixtures with limited labeling required. Past studies have illustrated how MALDI-TOF mass spectrometry is a practical tool for absolute and relative quantification of peptides and proteins. For example, using an internal standard of a similar molecular weight is sufficient for generation of a calibration curve.^33^ Quantitation of total uptake of L-CPPs was achieved using heavy atom-labeled internal standards.^34,35^ While this assay provided information regarding whole cell uptake of CPPs and CPP-peptide conjugates, it is limited by the need for heavy-atom labeling and the rapid degradation of L-peptides.^36^ A method for circumnavigating the need for spike-in of heavy atom-labeled standards was developed for the relative quantification of phosphopeptides.^37^ However, the proteolytic stability of D-peptides would facilitate their recovery and analysis as a mixture from inside cells and animals, allowing for the use of a new metric of antisense delivery efficiency.

Here we report that compared to the native L-forms, the mirror image forms of several sequences were equally able to deliver antisense molecules to the nucleus, but their increased proteolytic stability simplified mass spectrometry-based characterization following cytosolic delivery. Cytosolic delivery can be quantified based on the recovery of intact constructs from inside the cell. We profiled the uptake of biotinylated CPPs and PMO-CPPs to determine their relative concentrations in the whole cell and cytosol using careful extraction with digitonin and direct detection via MALDI-TOF. By comparing PMO delivery activity to relative internal concentration, we can derive a new metric for cargo delivery efficiency that may be useful for future development of CPPs for PMO delivery.

## Results and Discussion

### Mirror image peptides have same PMO delivery activity as native forms

We first established that several mirror image peptides could deliver a model PMO molecule to the nucleus of cells with similar efficacy to their native L-forms. We selected commonly studied, cationic peptides with precedented PMO delivery activity, but not known for their dependence on secondary structure or receptor interaction, and synthesized them in their L and D-form. The first iteration of these constructs contained a biotin linked through a 6-aminohexanoic acid residue and a trypsin-cleavable motif in between the peptide and the cargo. The cargo portion contained an azide for conjugation to PMO, and a biotin for use in affinity capture (Fig. 1A). The final constructs included cationic, oligoarginine, amphipathic, and hydrophobic sequences.

**Fig 1.**
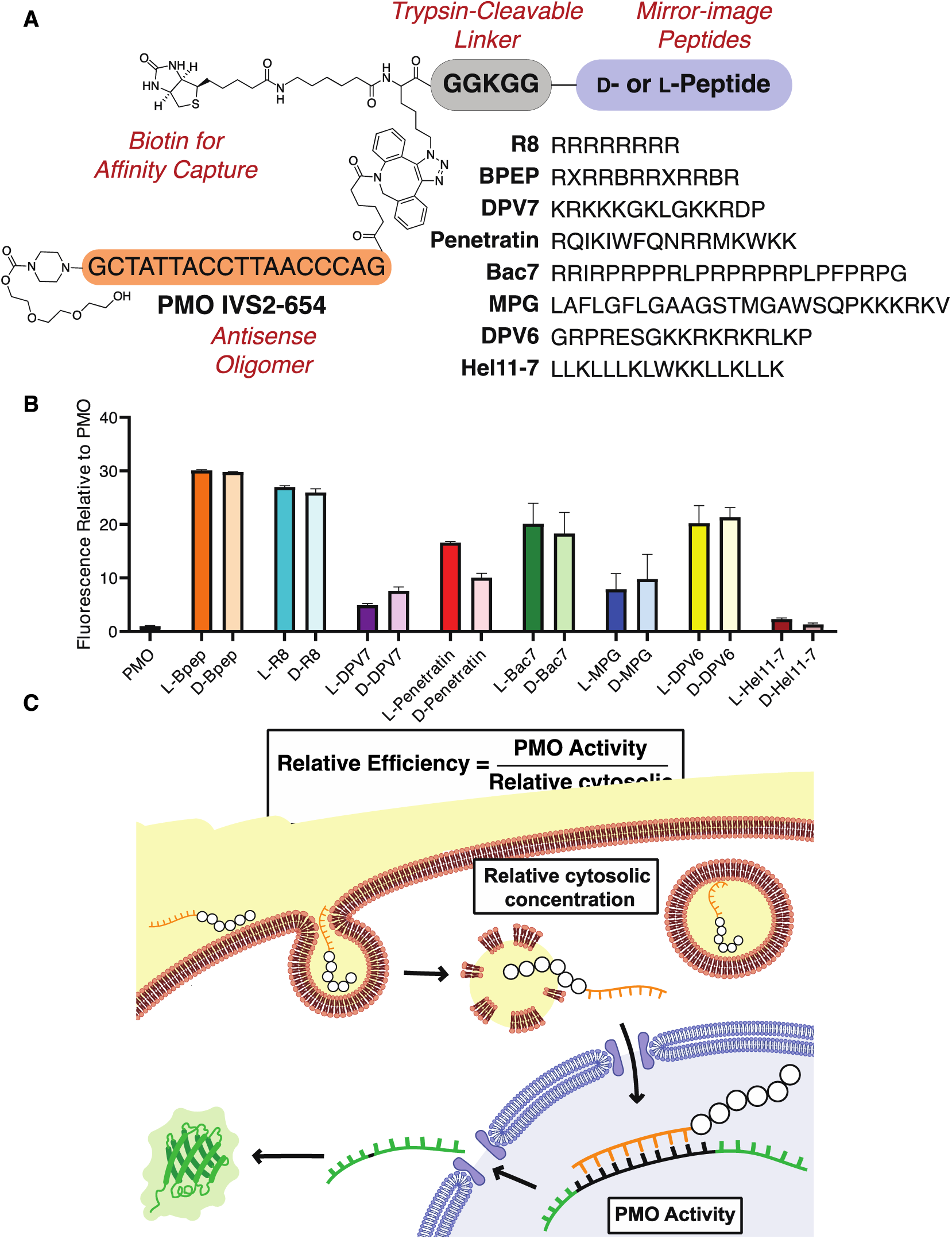
Mirror image cell-penetrating peptides have the similar PMO delivery activity as their native counterparts. (A) General construction of conjugates studied, including the ten studied CPP sequences. Macromolecular cargo PMO IVS2-654 is attached to the N-terminus of the peptides, along with a biotin handle for subsequent affinity capture. A trypsin-cleavable linker connects the cargo to the peptides. (B) Shown is activity data from the EGFP 654 assay conducted with D- and L-forms of several sequences at 5 µM. These initial constructs tested contained a 6-aminohexanoic acid moiety between the biotin and the peptide. Each bar represents group mean ± SD, N = 3 distinct samples from a single biological replicate. Experiment was repeated with similar results, shown in the SI (C) Schematic of how PMO-CPPs may enter the cell to perform exon-skipping activity. While activity assays give information of how much active PMO reached its target, a mass spectrometry-based assay may give information on the amount of PMO-CPP located in the cell and cytosol. Comparing these metrics provides a new estimate of CPP efficiency.

In total, fourteen PMO-peptide sequences were synthesized in their D- and L-form and tested in an activity-based in vitro assay.^22,23^ To synthesize the constructs, L-peptides were synthesized via automated fast-flow peptide synthesis, and D-peptides were synthesized using semi-automated fast flow peptide synthesis (SI). Azido-lysine and biotin moieties were added to the N-terminus of the peptides manually, and the peptides were simultaneously cleaved and deprotected before purification via RP-HPLC. PMO was modified with a dibenzocyclooctyne (DBCO) moiety and purified before attachment to the azido-peptides via strain-promoted azide-alkyne cycloaddition in water. Purified constructs were then tested using an activity-based readout in which nuclear delivery results in fluorescence. Briefly, HeLa cells stably transfected with an EGFP gene interrupted by a mutated intron of ß-globin (IVS2-654) produce a non-fluorescent EGFP protein. Successful delivery of PMO IVS2-654 to the nucleus results in corrective splicing and EGFP synthesis. The amount of PMO delivered to the nucleus is therefore correlated with EGFP fluorescence, quantified by flow cytometry. Activity is reported as mean fluorescence intensity (MFI) relative to PMO alone. This activity assay provides indirect information on how much active PMO is delivered to the nucleus. Relative efficiency of a PMO-CPP could be characterized by comparing activity to internal concentration, as discussed later (Fig. 1C).

Dose-response studies with sequences in the D- and L-form confirmed similar activities between mirror image CPPs. From our initial proof-of-concept experiment, involving eight sequences in L- and D-form tested at a single concentration (5 µM), there was not a significant difference between the activities of the mirror image peptides (Fig 1B). PMO-D-penetratin and PMO-D/L-Hel11-7 demonstrated some toxicity and were discontinued from study (SI). Constructs in subsequent experiments did not contain the 6-aminohexanoic acid linker, and also contained a single Trp residue for determining their concentration by UV-Vis. These constructs were tested at varying concentrations in the EGFP 654 assay, and the results further suggested that mirror image peptides shared nearly identical PMO delivery activities (Fig. 2A, SI). Though striking similarities were observed for the selected peptides, this similarity is certainly not expected for all CPPs and cargoes, including those that rely on secondary structure or receptor-mediated uptake. Furthermore, the supernatant from these assays was tested for lactate dehydrogenase (LDH) release, indicative of membrane toxicity, and it was confirmed that the constructs did not elicit membrane toxicity at the doses tested (SI).

**Fig 2.**
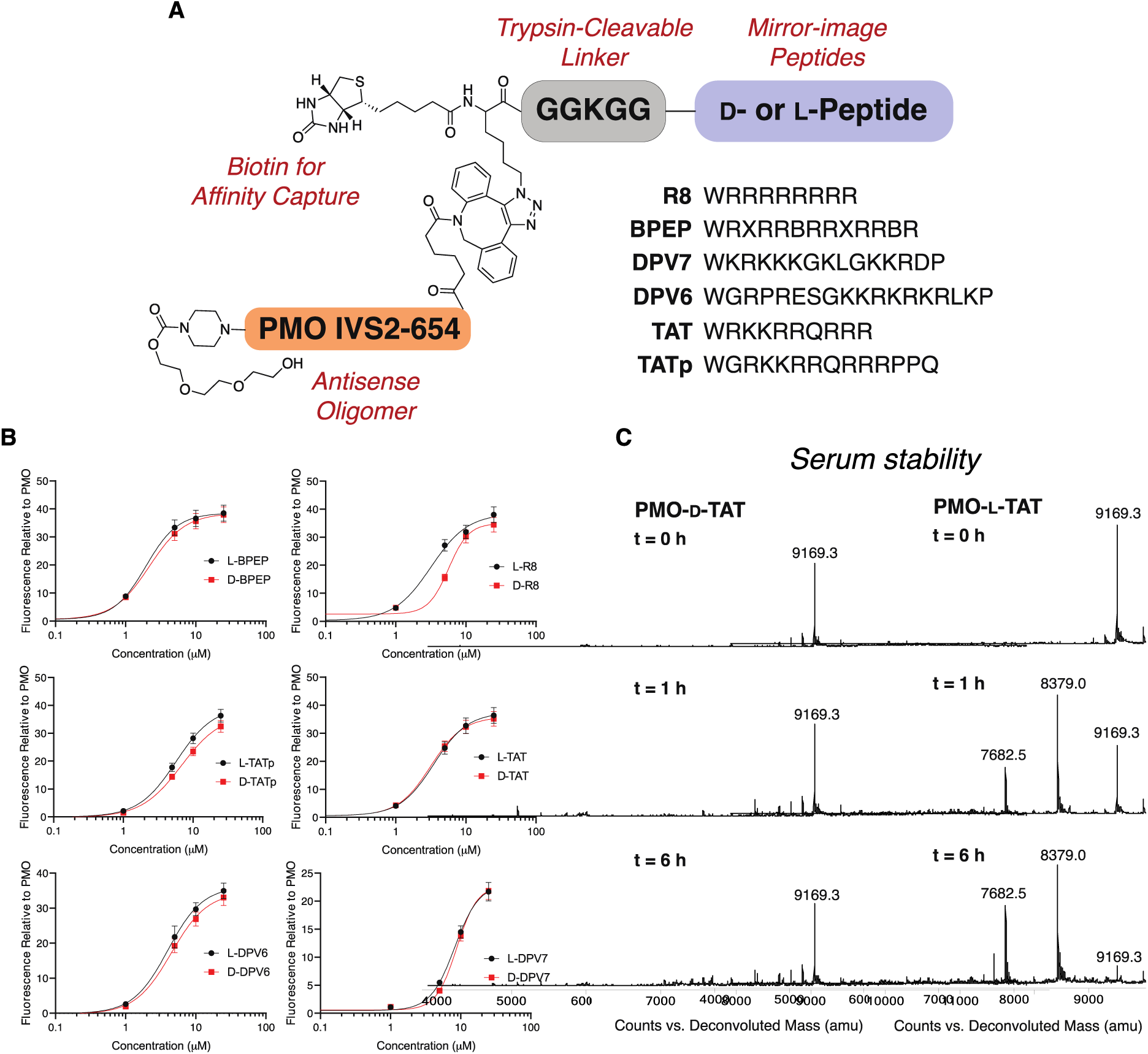
Mirror image cell-penetrating peptides have the same PMO delivery activity as their native counterparts while remaining proteolytically stable. (A) Construction and sequences of PMO-peptides used in subsequent experiments. The aminohexanoic acid linker was removed, and each peptide contains a single tryptophan residue for quantitation by UV-Vis. (B) Dose-response curves of D- and L-forms of several sequences at varying concentrations. No membrane toxicity was observed after analyzing the supernatant of these experiments (SI). Each point represents group mean ± SD, N = 3 distinct samples from a representative biological replicate. This experiment was repeated twice independently with similar results (SI) (C) Mass spectra of PMO-D- and L-conjugates following incubation with human serum. The L-variant is degraded within 1 h, while the D-variant is stable after 6 h. Spectra from other PMO-CPPs in shown in Supplemental section 2.4.

### Mirror image peptides are proteolytically stable

Our primary motivation for investigating mirror image peptides for transporting PMO was that the D-form would be stable against proteolysis and thus would match this property of the PMO cargo. D-peptides are indeed stable against degradation, illustrated by a time-course study in which both forms of PMO-CPPs were incubated in 25% human serum. While the studied PMO-D-CPPs remained intact 24 h later, the L-forms rapidly degraded into multiple fragments, leaving the parent construct as a minor product after only one hour of incubation (Fig 2B). This observation furthers the notion that L-peptides are not suitable for investigation using mass spectrometry after recovery from a biological setting. However, D-peptide conjugates can be recovered from a biological environment such as serum without suffering degradation, simplifying their characterization via mass-spectrometry.

### Mirror image PMO-peptides can be recovered from inside cells

The proteolytic stability of D-peptides simplifies the recovery of mixtures of intact constructs after being internalized into cells. While MALDI-ToF has been used previously to analyze quantity of L-peptides recovered from inside cells, it has not yet been used to profile drug-peptide conjugates, D-peptides, or mixtures of more than three conjugates at a time.^34,35^ The use of D-peptides would facilitate the analysis of a mixture of conjugates, because without degradation, only the parent peak would be observed. Moreover, this platform has not yet been used to study peptides recovered from sub-cellular fractions. Through a series of experiments, we show that mixtures of intact PMO-D-CPPs can be recovered from the cytosol of cells and analyzed by MALDI-ToF to estimate their relative abundances without the need for isotope labeling or standard curves (Fig 3A).

**Fig 3.**
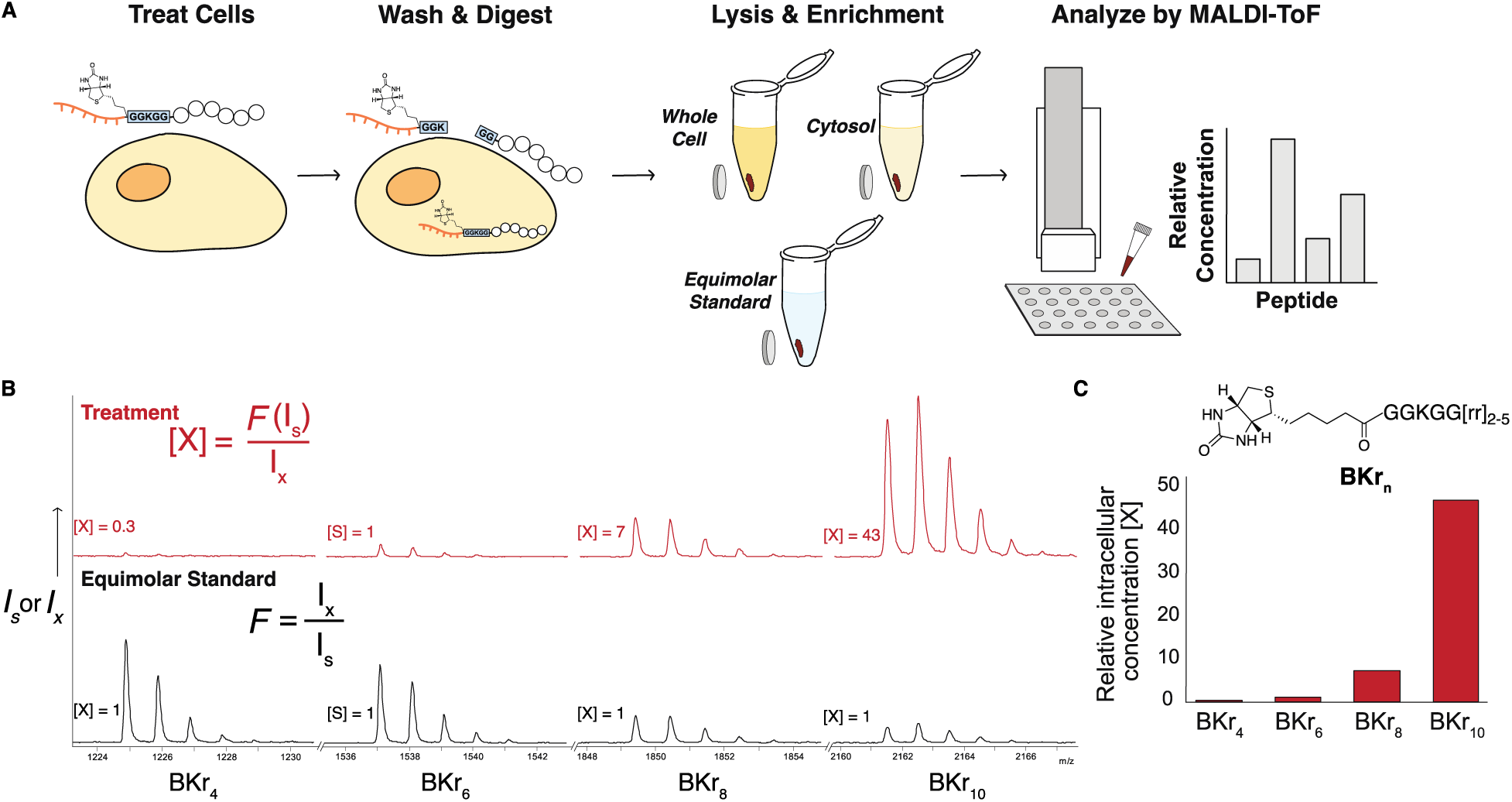
Uptake assay reveals relative concentrations of intact construct inside the cell. (A) Workflow of the uptake assay; cells are treated with PMO-D-CPPs, washed, and lysed to extract the whole cell lysate or the cytosol. A trypsin-cleavable L-linker ensures extracellular constructs are not recovered. Constructs are immobilized on magnetic streptavidin beads, washed, and plated directly for MALDI analysis. Intensities of analytes compared to an equimolar standard provide relative concentrations. (B) MALDI-TOF mass spectra displaying ions of intact biotinylated D-polyarginine peptides, isolated after internalization into HeLa cells. Spectra show ions corresponding to intact conjugates in the equimolar spike-in (black) and the whole cell lysate of cells treated with equimolar mixture (red). (C) By comparing relative intensities in the equimolar standard, the response factor (*F*) was determined and used to calculate the fold change in concentration in the experimental samples, shown as bar graph, normalized to BKr_4_. Also shown is the equation used to determine relative concentration: *I* (intensity), *[X]* (sample concentration), *F* (response factor), *X* (sample), *S* (standard).

Our first step was to recapitulate a known empirical trend that more Arg residues leads to greater uptake. We began our assay with a simple model system of four polyarginine peptides with a trypsin cleavable linker and a single biotin label. HeLa cells were incubated with biotin-K-D-Arg_4_, D-Arg_6,_ D-Arg_8,_ and D-Arg_10_ for 1 h. The cells were then washed extensively with PBS and heparin and trypsinized to lift the cells as well as to cleave the L-linker on the constructs, preventing recovery of extracellular peptides. The whole cell lysate was then prepared using RIPA buffer. Fully intact biotinylated peptides were captured with magnetic streptavidin Dynabeads, washed, and plated directly for MALDI analysis. Also plated were Dynabeads incubated in an equimolar mixture of the same constructs as determined by UV-Vis (Fig 3B).

The relative concentration of peptides on the beads can be estimated by determining the analyte’s response factor (F) from the equimolar standard (Fig 3C). In the standard, each analyte’s concentration is 1 mM, and each analyte’s response factor (F) is determined by normalizing their intensities to an internal standard (S). Here, BKr_6_ was selected as the arbitrary standard, where F = 1. The response factor of each analyte should remain consistent across samples that contain the same analytes^33,37^, and was used to calculate the fold change in concentration in the experimental samples. The relative concentrations [X] of the analytes, normalized to the ‘internal standard’ BKr_6_ are shown as a bar graph (Fig 3C). There is a clear increase in concentration of the constructs with more Arg residues, with BKr_10_ having 40-fold greater concentration than BKr_6_. This trend of greater number of Arg residues leading to greater uptake is already well documented in the literature.^38,39^

### PMO-D-CPPs can be extracted from cytosol and analyzed by MADI

Next, we found that these PMO-CPP constructs likely enter via energy-dependent endocytosis. When evaluating CPP delivery efficiency, considering the mechanism of uptake and potential for endosomal entrapment is necessary. In contrast with CPPs alone which can enter via passive diffusion,^38,40^ we hypothesized that the uptake mechanism of these constructs was likely endocytosis, considering the size of the PMO cargo as well as previous studies that we have conducted on similar conjugates.^22,23^ Using a panel of chemical endocytosis inhibitors, we performed a pulse-chase EGFP 654 assay format in which cells were pre-incubated with inhibitors before treatment with the L- and D-forms of PMO-DPV7 and PMO-Bac7 (Supplementary Information section 2.3). Analysis by flow cytometry revealed that chlorpromazine reduced activity in a dose-dependent manner (Fig 4A). Chlorpromazine is an inhibitor of clathrin-mediated endocytosis and has previously been observed to inhibit activity of similar PMO constructs. While it is possible that multiple uptake mechanisms are occurring, these PMO-CPP conjugates are likely taken up by active transport.

**Fig. 4.**
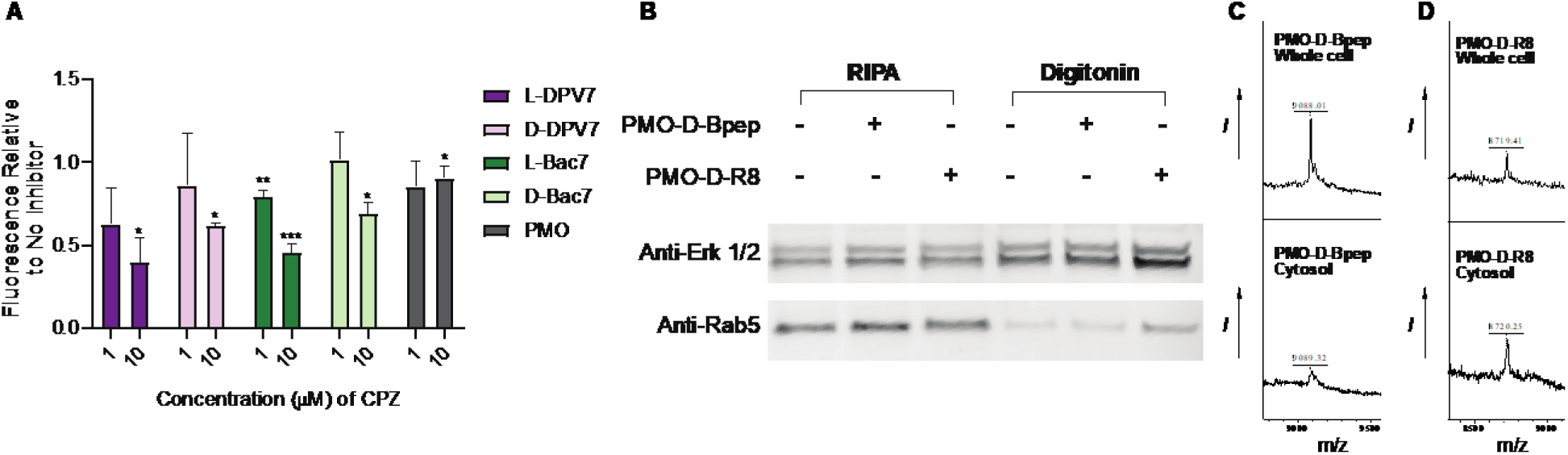
PMO-D-CPPs enter via endocytosis, and can be detected in whole cell and cytosolic lysate by MALDI-TOF. (A) Shown are PMO activities from an EGFP 654 assay in which cells are treated in a pulse-chase format with chemical endocytosis inhibitors followed by PMO-conjugates at 5 µM. Chlorpromazine (CPZ) produces a dose-dependent inhibition of PMO activity. Bars represent group mean ± SD, N = 3 distinct samples from a single biological replicate. (B) Western blot demonstrating extraction of whole cell lysate and cytosolic fraction with RIPA and digitonin buffer, respectively. Erk 1/2 is a cytosolic marker, whereas Rab5 is a late endosomal marker. (C) and (D) show example MALDI spectra following uptake analysis of lysates from (B) containing PMO-D-Bpep and PMO-D-R8, respectively. Intact construct was detected in the whole cell (top) as well as cytosolic (bottom) fractions.

Knowing that PMO-CPPs enter via endocytosis, we assert that extracting the cytosol and comparing to whole cell lysate is critical when evaluating relative concentrations of constructs internalized into cells. Not all endocytosed compounds are able to escape the endosome, and endosomal entrapment would lead to less active PMO delivered into the cytosol, measured by a lower cytosolic concentration relative to whole-cell lysate. Therefore, we then extracted biotinylated PMO-CPPs from the cytosol as well as the whole cell lysate following internalization and detected them by Western blot and MALDI. Individually, we incubated PMO-D-R8 or PMO-D-Bpep at 5 µM with HeLa cells in a 12-well plate for 1 h before washing with heparin and digesting with trypsin. The cytosol was extracted using Digitonin buffer, which selectively permeabilizes the outer membrane. RIPA buffer was used to prepare whole cell lysates. To confirm cytosolic extraction, a portion of each sample was analyzed via Western blot using a cytosolic marker (Erk 1/2) and a late-endosomal marker (Rab5). Samples of cytosolic extract have markedly reduced Rab5 while all samples contain Erk 1/2 (Fig 4B). Finally, as with the biotinylated peptides, the PMO-CPPs were then extracted from the samples with Streptavidin coated magnetic Dynabeads, washed extensively, and analyzed via MALDI-TOF. We indeed detected PMO-D-R8 and PMO-D-Bpep in their respective samples, presenting the first instance of an intact peptide-oligonucleotide conjugate being extracted from cells and analyzed by mass spectrometry (Fig 4C-D). Moreover, we found that incubation at lower temperature appeared to inhibit cytosolic localization of PMO-D-CPPs, but resulted in equal relative concentrations between whole cell and cytosolic fractions for biotin-D-CPPs (Supplementary Information section 3.2).

Next, we expanded the analytes tested and profiled the uptake of six biotin-D-CPPs in the cytosol of distinct cell lines. Biotin-D-R8, TAT, Bpep, DPV7, TATp, and DPV6 were profiled by MALDI in both HeLa (Fig 5A) and C2C12 mouse myoblast (Fig 5B) cell lines. We observed different uptake patterns between the two cell lines; polyarginine was significantly more abundant in the C2C12 cells compared to the other peptides, and DPV6 and DPV7 were not detected in the cytosol. In HeLa, polyarginine had the highest relative concentration in both cytosol and whole cell, and DPV7 was again not detected in the cytosol. However, TAT, TATp, and DPV6 all had similar relative cytosolic concentrations. With these experiments we demonstrated that this profiling platform could determine the relative concentration of six intact peptides extracted from whole cell and cytosol of two distinct cell lines.

**Fig. 5.**
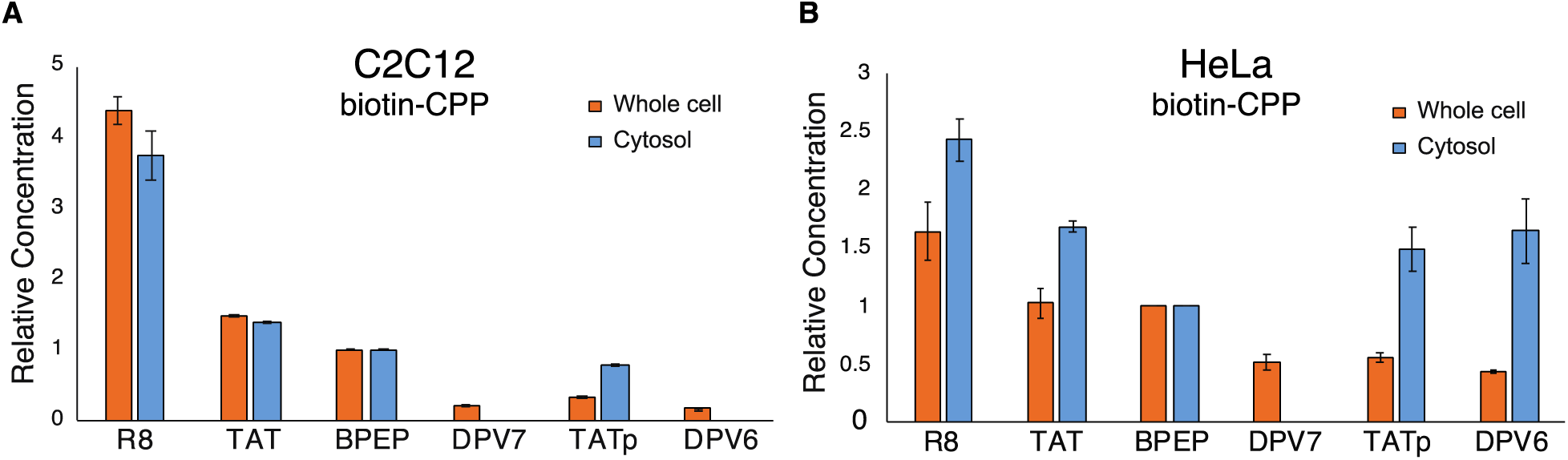
Uptake of biotin-CPPs can be profiled in different cell lines. (A) Bar graph showing concentrations of biotin-CPPs relative to BPEP in the whole cell and cytosolic extracts of (A) C2C12 mouse myoblast cells and (B) HeLa cells following 1 h treatment at 37 °C. Bars represent group mean +- SD N = 2 replicate samples. (A-B) Relative concentration is normalized to TATp. (C-D) Relative concentration is normalized to Bpep.

Finally, we profiled uptake of six PMO-D-CPPs. PMO-D-R8, TAT, BPEP, DPV7, TATp, and DPV6 were profiled in HeLa cells as usual. Extraction of cytosol was confirmed by Western blot (SI) and the samples were analyzed by MALDI. Relative concentrations were normalized to BPEP. In the whole cell extract, the relative concentrations were generally consistent across peptides, with DPV6 having 1.5-fold higher relative concentration. On the other hand, relative concentrations in the cytosol were lower for the polyarginine peptides R8 and TAT, and highest for BPEP and DPV6 (Fig 6A). By comparing the relative concentrations in cytosolic and whole-cell lysates, the relative efficiencies demonstrate an effective metric for determining which of the six CPPs can effectively deliver PMO into the cytosol (Fig. 6B).

**Fig. 6.**
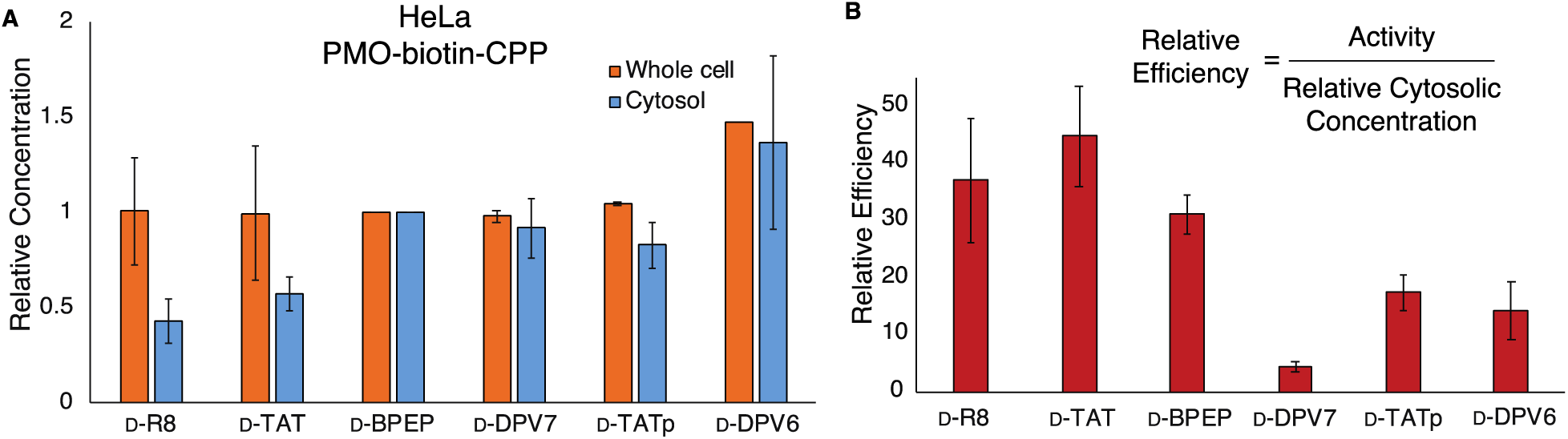
Mass spectrometry-based profiling combined with activity gives new efficiency metric for PMO-CPPs. (A) Bar graph showing concentrations of PMO-biotin-CPPs relative to BPEP in the whole cell and cytosolic extracts of Hela cells. Relative concentration is normalized to BPEP. (B) Bar graph showing relative efficiency (PMO activity / relative cytosolic concentration) of PMO-CPPs. Bars show group mean ± SD, N = 2 distinct samples from a single biological replicate, except for the whole cell condition of (A) in which N = 2 distinct samples from two independent biological replicates.

## Conclusion

Here we show that the all-D version of several known cell-penetrating peptides can deliver an antisense oligonucleotide to the nucleus of cells at the same efficiency as their native L-form. While the respective activities were nearly identical in cells, the D-forms were resistant to serum proteolysis. This stability enabled recovery and mass spectrometric analysis of the D-peptides from cytosolic and whole cell lysates. By comparing PMO activity to relative internalized concentrations, we obtained a metric for PMO delivery efficiency.

While it would not be expected that the mirror image of all CPPs would retain delivery activity, the striking similarities observed for the mirror image peptides selected in this study suggests that D-peptides should be studied further. Despite similar activity in cells, the difference in proteolytic stability indicates the potential for differential activities in animals. Proteolytic degradation of peptide therapeutics has been considered a major weakness limiting therapeutic investigation.^41^ However, mutations to D-amino acids, or the study of entirely D-peptides, has the potential to improve pharmacokinetic properties of proteinogenic therapies. Avoidance of cleavage sites can enhance proteolytic stability and half-life in vivo, and can be achieved by integration of unnatural amino acids, including D-amino acids.^42^ An all-D polyproline CPP was shown to be efficacious in mice^18^, and PMO-D-CPPs were previously found to be the most stable among tested B-peptide analogs.^15^ Despite these promising findings, the activities of PMO-D-CPPs had not previously been explored. We found that in HeLa cells, several CPP sequences had nearly identical delivery activities in their D- and L-forms, while the D-form remained completely proteolytically stable.

Besides potential in vivo applications, the proteostability of D-peptides also simplified their direct analysis by mass spectrometry following recovery from biological milieu. Previous reports demonstrated that biotinylated L-peptides could be recovered from inside whole cell lysate and quantified by MALDI-ToF, and cleavage products were identified and accounted for.^7,34^ Here, by focusing only on fully intact constructs, we analyzed a mixture of six individual biotinylated D-peptides from inside cells by observing only the intact parent ions. Moreover, we demonstrated relative quantification of a mixture of biotinylated peptides recovered from cytosolic extract. Analysis of the cytosolic portion is critical when studying and developing cell-penetrating peptides, considering that whole cell uptake does not correlate with cytosolic delivery due to the possibility of endosomal entrapment.

Finally, we showed relative quantification of intact biotinylated PMO-D-CPPs in whole cell lysate. By combining relative internal concentrations with PMO delivery activity, we obtained a metric for relative delivery efficiency. Delivery efficiency would be a useful metric for comparing CPPs delivering active therapeutic cargo as it takes into account both the activity of the cargo as well as the internal concentration. A highly efficient peptide would have high activity with low internal concentration, and thus a high relative efficiency. Using these metrics when investigating CPPs would be useful in narrowing the scope of sequences early in development to exclude sequences which accumulate in the endosomes or otherwise do not efficiently deliver active cargo.

Exploration of mirror image CPPs would enter into a largely untapped chemical space. We recently described a machine learning-based platform for the discovery of nuclear-targeting peptides containing unnatural amino acids.^23^ While the unnatural residues included in this work were non-alpha-amino acids, future applications of this method could include D-amino acid substitutions, or fully mirror image CPPs in order to discover highly active, novel sequences that are completely stable.

## Experimental Section

### Preparation of PMO-Peptides

All peptides were prepared via fast-flow peptide synthesis, detailed fully in the Supporting Information. Briefly, peptides were synthesized on a 0.1 mmol scale using an automated fast-flow peptide synthesizer for L-peptides and a semi-automated fast-flow peptide synthesizer for D-peptides. Automated^43^ and semi-automated^44^ synthesis conditions were used as previously reported. To couple unnatural amino acids or to cap the peptide (e.g. with 4-pentynoic acid), the resin was incubated for 30 min at room temperature with amino acid (1 mmol) dissolved in 2.5 mL 0.4 M HATU in DMF with 500 μL diisopropylethylamine. After completion of the synthesis, the resin was washed 3 times with dichloromethane and dried under vacuum.

Each peptide was subjected to simultaneous global side-chain deprotection and cleavage from resin by treatment with 5 mL of 94% trifluoroacetic acid (TFA), 2.5% thioanisole, 2.5% water, and 1% triisopropylsilane (TIPS) (v/v) at room temperature for 2 to 4 hours. Cleaved peptide was isolated by trituration in cold ether, dissolved in 50% aqueous acetonitrile 0.1% TFA and lyophilized before purification via mass-directed semi-preparative reversed-phase HPLC.

PMO-DBCO (1 eq, 5 mM, water) was conjugated to azido-peptides (2 eq, 5 mM, water) at room temperature for 2 h. Reaction progress was monitored by LCMS and purified when PMO-DBCO was consumed. Purification was conducted using mass-directed HPLC (Solvent A: 100 mM ammonium acetate in water, Solvent B: acetonitrile).

### EGFP Assay

HeLa 654 cells were maintained in MEM supplemented with 10% (v/v) fetal bovine serum (FBS) and 1% (v/v) penicillin-streptomycin at 37 °C and 5% CO_2_. 18 h prior to treatment, the cells were plated at a density of 5,000 cells per well in a 96-well plate in MEM supplemented with 10% FBS and 1% penicillin-streptomycin.

PMO-peptides were dissolved in cation-free PBS at a concentration of 1 mM (determined by UV) before being diluted in MEM. Cells were incubated at the designated concentrations for 22 h at 37 °C and 5% CO_2._ The treatment media was removed, and the cells were washed once before being incubated with 0.25 % Trypsin-EDTA for 15 min at 37 °C and 5% CO_2_. Lifted cells were transferred to a V-bottom 96-well plate and washed once with PBS, before being resuspended in PBS containing 2% FBS and 2 µg/mL propidium iodide (PI). Flow cytometry analysis was carried out on a BD LSRII flow cytometer at the Koch Institute. Gates were applied to the data to ensure that cells that were positive for propidium iodide or had forward/side scatter readings that were sufficiently different from the main cell population were excluded. Each sample was capped at 5,000 gated events.

Analysis was conducted using Graphpad Prism 7 and FlowJo. For each sample, the mean fluorescence intensity (MFI) and the number of gated cells was measured. To report activity, triplicate MFI values were averaged and normalized to the PMO alone condition.

### Uptake Assay

The uptake assay was adapted from a previously reported protocol and is detailed fully in the Supporting Information.^35^ Briefly, cells were plated in a 12-well plate the evening before the experiment. On the day of, cells were treated at varying concentrations of PMO-biotin-peptide or biotin-peptide at varying temperatures and durations. Treatment media was removed, and cells were washed with fresh media before incubated with porcine heparin for 5 min to dissociate membrane-bound peptide. Cells were lifted and membrane-bound peptide was cleaved by incubation in trypsin for 5 min. Collected cells were pelleted and washed before being digested with RIPA buffer (for whole cell extract) or digitonin buffer (for cytosolic extract) on ice. Supernatant was collected and protein concentration was quantified using a BCA protein assay kit. A portion of some samples were retained for analysis by Western blot. The remaining supernatant was incubated with magnetic streptavidin Dynabeads overnight at 4 °C.

Dynabeads were washed with a series of buffers: 2 × 100 µL Buffer A (50 mM Tris-HCl (pH 7.4) and 0.1 mg/mL BSA), 2 × 100 µL Buffer B (50 mM Tris-HCl (pH 7.4), 0.1 mg/mL BSA, and 0.1% SDS), 2 × 100 µL Buffer C (50 mM Tris-HCl (pH 7.4), 0.1 mg/mL BSA, and 1 M NaCl), and 2 × 100 µL water. Washed beads were then resuspended in CHCA MALDI matrix and plated directly on a MALDI plate. The plate was analyzed using MALDI-ToF on a high-resolution Bruker Autoflex LRF Speed mass spectrometer in linear positive mode.

Relative concentrations of peptides in the mixture were determined as follows. Analytes in a mixture ionize according to their response factor (F). F was determined by normalizing the intensities of each analyte to one analyte in the control sample, where the concentration of each analyte is arbitrarily set to 1. The values of F is then used in the experimental spectra containing the same mixture of analytes to determine their relative concentrations.

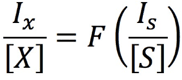

### Statistics

Statistical analysis and graphing was performed using Prism (Graphpad) or Excel (Microsoft). Concentration-response curves were fitted using Prism using nonlinear regression. The listed replicates for each experiment indicates the number of distinct samples measured for a given assay. Significance for activities between constructs was determined using a student’s two-sided, unpaired t-test.

## Supporting Information

Supporting information is available free of charge via the Internet. Materials, methods for peptide preparation, LC-MS analysis (Section 1); activity and toxicity assays, serum stability assay (Section 2); methods for uptake assay, additional MALDI-ToF experiments (Section 3); Gel images (Appendix I); and LC-MS Characterization (Appendix II).

## Data and materials availability

All data is available in the main text or the Supporting Information.

## Acknowledgements

This research was funded by Sarepta Therapeutics. C.K.S. (4000057398) and C.E.F. (4000057441) acknowledge the National Science Foundation Graduate Research Fellowship (NSF Grant No. 1122374) for research support. We thank the Swanson Biotechnology Center Flow Cytometry Facility at the Koch Institute for the use of their flow cytometers, and MIT’s Department of Chemistry Instrumentation Facility for the use of the MALDI-ToF.

