## Supporting Information for "Cell-Penetrating D-Peptides Retain Antisense Morpholino Oligomer Delivery Activity"

### 1 Materials and General Methods

#### 1.1 Reagents and Solvents

H-Rink Amide-ChemMatrix resin was obtained from PCAS BioMatrix Inc. (St-Jean-sur-Richelieu, Quebec, Canada). 1-[Bis(dimethylamino)methylene]-1H-1,2,3-triazolo[4,5-b]pyridinium-3- oxid-hexafluorophosphate (HATU), 4-pentynoic acid, 5-azidopentanoic acid, Fmoc-β-Ala-OH, Fmoc-6-aminohexanoic acid, and Fmoc-L-Lys($N_3$) were purchased from Chem-Impex International (Wood Dale, IL). PyAOP was purchased from P3 BioSystems (Louisville, KY). Fmoc-protected L-amino acids (Fmoc-Ala-OHx$H_2O$, Fmoc-Arg(Pbf)-OH; Fmoc-Asn(Trt)-OH; Fmoc-Asp-(O*t*-Bu)-OH; Fmoc-Gln(Trt)-OH; Fmoc-Glu(O*t*-Bu)-OH; Fmoc-Gly-OH; Fmoc-His(Trt)-OH; Fmoc-Ile-OH; Fmoc-Leu-OH; Fmoc-Lys(Boc)-OH; Fmoc-Met-OH; Fmoc-Phe-OH; Fmoc-Pro-OH; Fmoc-Ser(But)-OH; Fmoc-Thr(*t*-Bu)-OH; Fmoc-Trp(Boc)-OH; Fmoc-Tyr(*t*-Bu)-OH; Fmoc-Val-OH), were purchased from the Novabiochem-line from Sigma Millipore. Fmoc-protected D-amino acids were purchased from Chem Impex (Wood Dale, IL). Peptide synthesis-grade *N,N*-dimethylformamide (DMF), $CH_2Cl_2$, diethyl ether, and HPLC-grade acetonitrile were obtained from VWR International (Radnor, PA). All other reagents were purchased from Sigma-Aldrich (St. Louis, MO). Milli-Q water was used exclusively.

#### 1.2 Liquid-chromatography mass-spectrometry

LCMS analyses were performed on either an Agilent 6520 or 6545 Accurate-Mass Q-TOF LCMS (abbreviated as 6520 or 6545) coupled to an Agilent 1260 Infinity HPLC system, or an Agilent 6550 iFunnel Q-TOF LCMS system (abbreviated as 6550) coupled to an Agilent 1290 Infinity HPLC system. Mobile phases were: 0.1% formic acid in water (solvent A) and 0.1% formic acid in acetonitrile (solvent B). The following LCMS methods were used for characterization:

**Method A: 1-61% B over 9 min, Zorbax C3 column (6520 and 6545)**

<u>LC</u>: Zorbax 300SB-C3 column: 2.1 × 150 mm, 5 µm, column temperature: 40 °C, gradient: 0-2 min 1% B, 2-11 min 1-61% B, 11-12 min 61-95% B, 12-15 min 95% B; flow rate: 0.8 mL/min.

<u>MS</u>: Positive electrospray ionization (ESI) extended dynamic range mode in mass range 300–3000 *m*/*z*. MS is on from 4 to 11 min.

**Method B: 1-91% B over 9 min, Zorbax C18 column (6520 and 6545)**

LC: Zorbax 300SB-C3 column: 2.1 × 150 mm, 5 μm, column temperature: 40 °C, gradient: 0-2 min 1% B, 2-11 min 1-91% B, 11-12 min 91-95% B, 12-15 min 95% B; flow rate: 0.8 mL/min.
MS: Positive electrospray ionization (ESI) extended dynamic range mode in mass range 300–3000 *m*/*z*. MS is on from 4 to 11 min.

**Method C: 1-61% B over 10 min, Phenomenex Jupiter C4 column (6550)**

LC: Phenomenex Jupiter C4 column: 1.0 × 150 mm, 5 μm, column temperature: 40 °C, gradient: 0-2 min 1% B, 2-12 min 1-61% B, 12-16 min 61-90% B; 16-20 min 90% B; flow rate: 0.1 mL/min.
MS: Positive electrospray ionization (ESI) extended dynamic range mode in mass range 100–1700 *m*/*z*. MS is on from 4 to 12 min.

**Method D: 1-61% B over 10 min, Agilent EclipsePlus C18 column (6550)**

LC: Agilent EclipsePlus C18 RRHD column: 2.1 × 50 mm, 1.8 μm, column temperature: 40 °C, gradient: 0-2 min 1% B, 2-12 min, 1-61% B, 12-13 min, 61% B, 13-16 min, 1% B; flow rate: 0.1 mL/min.
MS: Positive electrospray ionization (ESI) extended dynamic range mode in mass range 300–3000 m/z. MS is on from 4 to 12 min. This method was used exclusively for characterization of the modular library.

All data were processed using Agilent MassHunter software package. Y-axis in all chromatograms shown represents total ion current (TIC) unless noted.

##### 1.3 General method for peptide preparation

Fast-flow Peptide Synthesis: Peptides were synthesized on a 0.1 mmol scale using an automated fast-flow peptide synthesizer for L-peptides and a semi-automated fast-flow peptide synthesizer for D-peptides. Automated synthesis conditions were used as previously reported.[1] Briefly, a 100 mg portion of ChemMatrix Rink Amide HYR resin was loaded into a reactor maintained at 90 ºC. All reagents were flowed at 40 mL/min with HPLC pumps through a stainless-steel loop maintained at 90 ºC before introduction into the reactor. For each coupling, 10 mL of a solution containing 0.4 M amino acid and 0.38 M HATU in DMF were mixed with 600 μL diisopropylethylamine and delivered to the reactor. Fmoc removal was accomplished using 10.4 mL of 20% (v/v) piperidine. Between each step, DMF (15 mL) was used to wash out the reactor. To couple unnatural amino acids or to cap the peptide (e.g. with 4-pentynoic acid), the resin was incubated for 30 min at room temperature with amino acid (1 mmol) dissolved in 2.5 mL 0.4 M

HATU in DMF with 500 μL diisopropylethylamine. After completion of the synthesis, the resin was washed 3 times with dichloromethane and dried under vacuum.

Semi-automated synthesis was carried out as previously described.[2] 1 mmol of amino acid was combined with 2.5 mL 0.4 M HATU and 500 μL DIEA and mixed before being delivered to the reactor containing resin via syringe pump at 6 mL/min. The reactor was submerged in a water bath heated to 70 ºC. An HPLC pump delivered either DMF (20 mL) for washing or 20 % piperidine/DMF (6.7 mL) for Fmoc deprotection, at 20 mL/min.

Peptide Cleavage and Deprotection: Each peptide was subjected to simultaneous global side-chain deprotection and cleavage from resin by treatment with 5 mL of 94% trifluoroacetic acid (TFA), 2.5% thioanisole, 2.5% water, and 1% triisopropylsilane (TIPS) (v/v) at room temperature for 2 to 4 hours. The cleavage cocktail was first concentrated by bubbling $N_2$ through the mixture, and cleaved peptide was precipitated and triturated with 40 mL of cold ether (chilled in dry ice). The crude product was pelleted by centrifugation for three minutes at 4,000 rpm and the ether was decanted. This wash step was repeated two more times. After the third wash, the pellet was dissolved in 50% water and 50% acetonitrile containing 0.1% TFA, filtered through a fritted syringe to remove the resin and lyophilized.

Peptide Purification: The peptides were dissolved in water and acetonitrile containing 0.1% TFA, filtered through a 0.22 μm nylon filter and purified by mass-directed semi-preparative reversed-phase HPLC. Solvent A was water with 0.1% TFA additive and Solvent B was acetonitrile with 0.1% TFA additive. A linear gradient that changed at a rate of 0.5% B/min was used. Most of the peptides were purified on an Agilent Zorbax SB C18 column: 9.4 x 250 mm, 5 μm. Using mass data about each fraction from the instrument, only pure fractions were pooled and lyophilized. The purity of the fraction pool was confirmed by LC-MS.

##### 1.4 PMO-DBCO Synthesis

PMO IVS-654 (50 mg, 8 μmol) was dissolved in 150 μL DMSO. To the solution was added a solution containing 2 equivalents of dibenzocyclooctyne acid (5.3 mg, 16 μmol) activated with HBTU (37.5μL of 0.4 M HBTU in DMF, 15 μmol) and DIEA (2.8 μL, 16 μmol) in 40 μL DMF (Final reaction volume = 0.23 mL). The reaction proceeded for 25 min before being quenched with 1 mL of water and 2 mL of ammonium hydroxide. The ammonium hydroxide hydrolyzed any ester formed during the course of the reaction. After 1 hour, the solution was diluted to 40 mL in water/acetonitrile and purified using reverse-phase HPLC (Agilent Zorbax SB C3 column: 21.2 x 100 mm, 5 µm) and a linear gradient from 2 to 60% B (solvent A: water; solvent B: acetonitrile) over 58 min (1% B / min). Using mass data about each fraction from the instrument, only pure fractions were pooled and lyophilized. The purity of the fraction pool was confirmed by LC-MS.

Conjugation to peptides

PMO-DBCO (1 eq, 5 mM, water) was conjugated to azido-peptides (1.5 eq, 5 mM, water) at room temperature for 2 h. Reaction progress was monitored by LCMS and purified when PMO-DBCO was consumed. Purification was conducted using mass-directed HPLC (Solvent A: 100 mM ammonium acetate in water, Solvent B: acetonitrile) with a linear gradient that changed at a rate of 0.5% B/min, on an Agilent Zorbax SB C13 column: 9.4 x 250 mm, 5 µm. Using mass data about each fraction from the instrument, only pure fractions were pooled and lyophilized. The purity of the fraction pool was confirmed by LC-MS.

### 2 Biological evaluation of peptides and PMO-peptides

#### 2.1 EGFP Assay

HeLa 654 cells obtained from the University of North Carolina Tissue Culture Core facility were maintained in MEM supplemented with 10% (v/v) fetal bovine serum (FBS) and 1% (v/v) penicillin-streptomycin at 37 °C and 5% $CO_2$. 18 h prior to treatment, the cells were plated at a density of 5,000 cells per well in a 96-well plate in MEM supplemented with 10% FBS and 1% penicillin-streptomycin.

For individual peptide testing, PMO-peptides were dissolved in PBS without $Ca^{2+}$ or $Mg^{2+}$ at a concentration of 1 mM (determined by UV) before being diluted in MEM. Cells were incubated at the designated concentrations for 22 h at 37 °C and 5% $CO_2$. Next, the treatment media was removed, and the cells were washed once before being incubated with 0.25 % Trypsin-EDTA for 15 min at 37 °C and 5% $CO_2$. Lifted cells were transferred to a V-bottom 96-well plate and washed once with PBS, before being resuspended in PBS containing 2% FBS and 2 µg/mL propidium iodide (PI). Flow cytometry analysis was carried out on a BD LSRII flow cytometer at the Koch Institute. Gates were applied to the data to ensure that cells that were positive for propidium iodide or had forward/side scatter readings that were sufficiently different from the main cell population were excluded. Each sample was capped at 5,000 gated events.

Analysis was conducted using Graphpad Prism 7 and FlowJo. For each sample, the mean fluorescence intensity (MFI) and the number of gated cells was measured. To report activity, triplicate MFI values were averaged and normalized to the PMO alone condition. For the final set of PMO-peptides evaluated, three biological replicates were performed. Individual biological replicates are shown below.

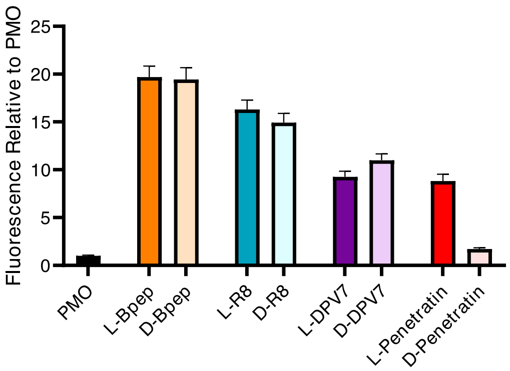

**Fig S1**. EGFP Assay of several L and D PMO-peptides with first generation linker. Performed at 5 µM. n = 3

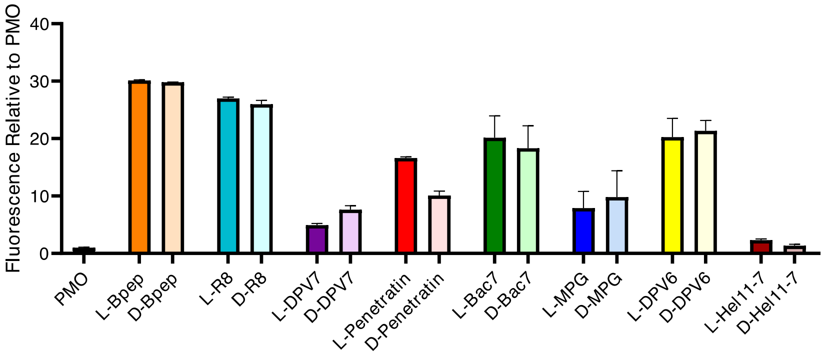

**Fig S2**. EGFP Assay of several L and D PMO-peptides with first generation linker. Performed at 5 µM. n = 3

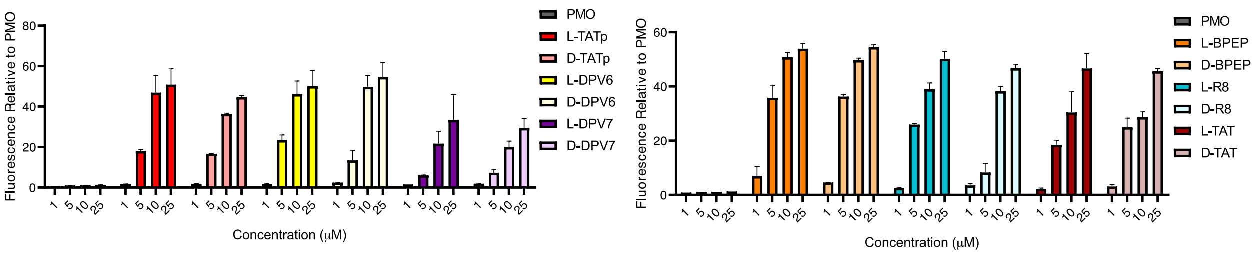

**Fig S3**. Single biological replicate of EGFP Assay of several L and D PMO-peptides with second generation linker. Performed at 5 µM. n = 3.

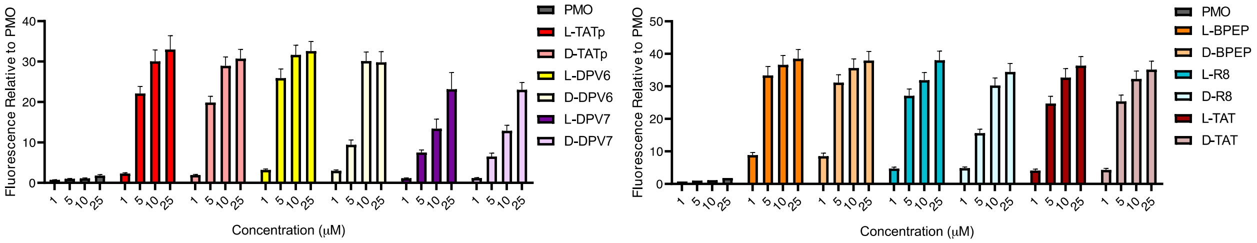

**Fig S4**. Single biological replicate of EGFP Assay of several L and D PMO-peptides with second generation linker. Performed at 5 μM. n = 3.

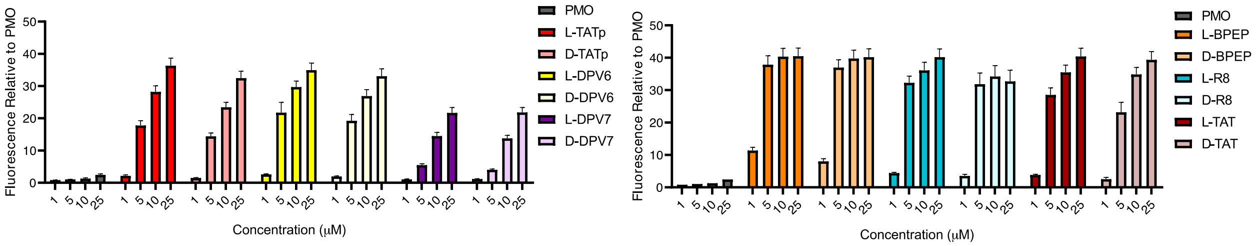

**Fig S5**. Single biological replicate of EGFP Assay of several L and D PMO-peptides with second generation linker. Performed at 5 μM. n = 3.

#### 2.2 Endocytosis Inhibition Assay

Chemical endocytosis inhibitors were used to probe the mechanism of delivery of PMO by these peptides in a pulse-chase format. We have conducted such analysis on similar PMO-peptide constructs previously with comparable outcomes.[3] For the PMO constructs, HeLa 654 cells were preincubated with various chemical inhibitors for 30 minutes before treatment with PMO-CPP constructs for three hours. The panel included: a panel of endocytosis inhibitors including: chlorpromazine (CPZ), which is demonstrated to interfere with clathrin-mediated endocytosis; cytochalasin D (CyD), which inhibits phagocytosis and micropinocytosis; wortmannin (Wrt), which alters various endocytosis pathways by inhibiting phosphatidylinositol kinases; EIPA (5-(N-ethyl-Nisopropyl) amiloride), which inhibits micropinocytosis; and Dynasore (Dyn), which also inhibits clathrin-mediated endocytosis.[4,5] Treatment media was then replaced with fresh media and the cells were incubated for 22 hours at 37 °C and 5% $CO_2$. Cells were then lifted as previously described and EGFP synthesis was measured by flow cytometry.

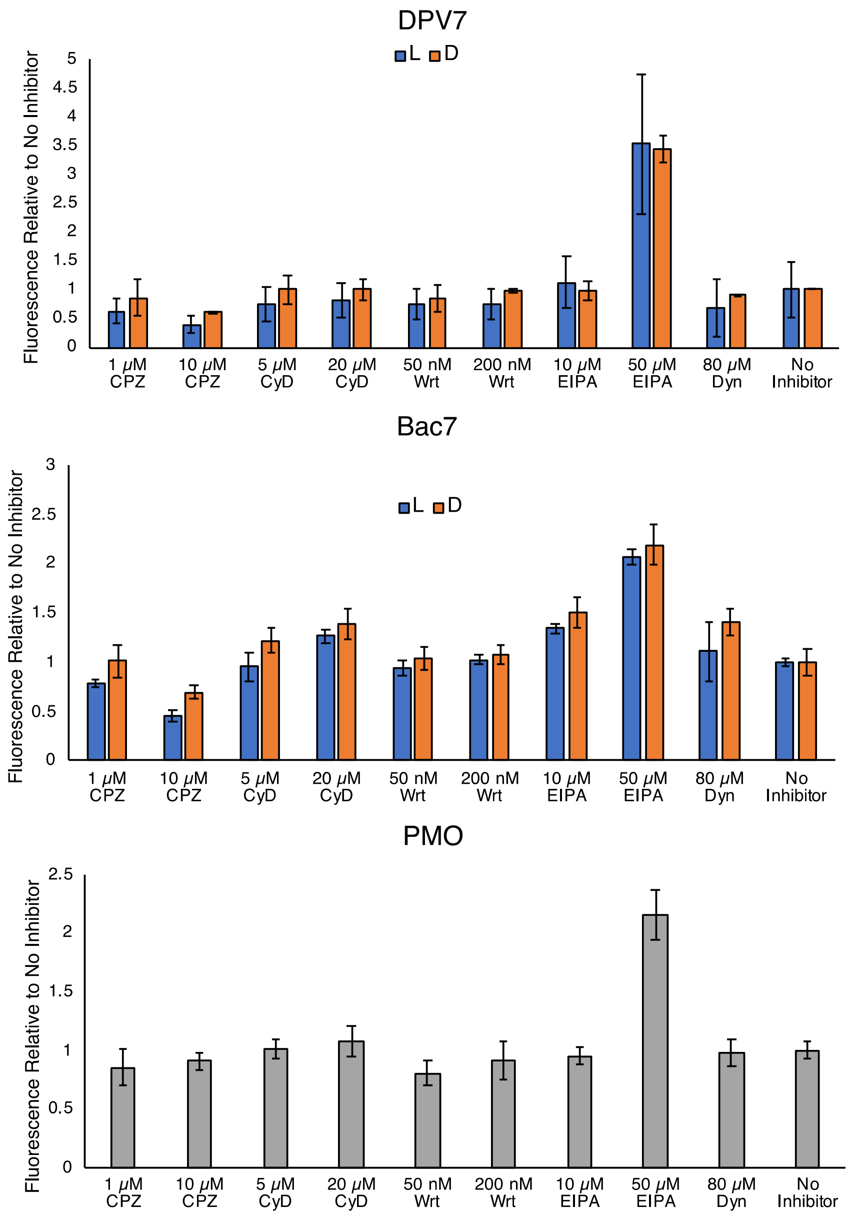

**Fig S6**. Pulse-chase EGFP Assay using several chemical endocytosis inhibitors, analyzing uptake pathways of PMO-Bac7, PMO-DPV7, and PMO alone. Concentration of PMO conjugate at 5 μM. n = 3.

#### 2.3 LDH Assay

Cytotoxicity assays were performed in HeLa 654 cells. Cell supernatant following treatment for flow cytometry was transferred to a new 96-well plate for analysis of LDH release. To each well of the 96-well plate containing supernatant was added CytoTox 96 Reagent (Promega). The plate was shielded from light and incubated at room temperature for 30 minutes. Equal volume of Stop

Solution was added to each well, mixed, and the absorbance of each well was measured at 490 nm. The blank measurement was subtracted from each measurement, and % LDH release was calculated as % cytotoxicity = 100 × Experimental LDH Release (OD490) / Maximum LDH Release (OD490).

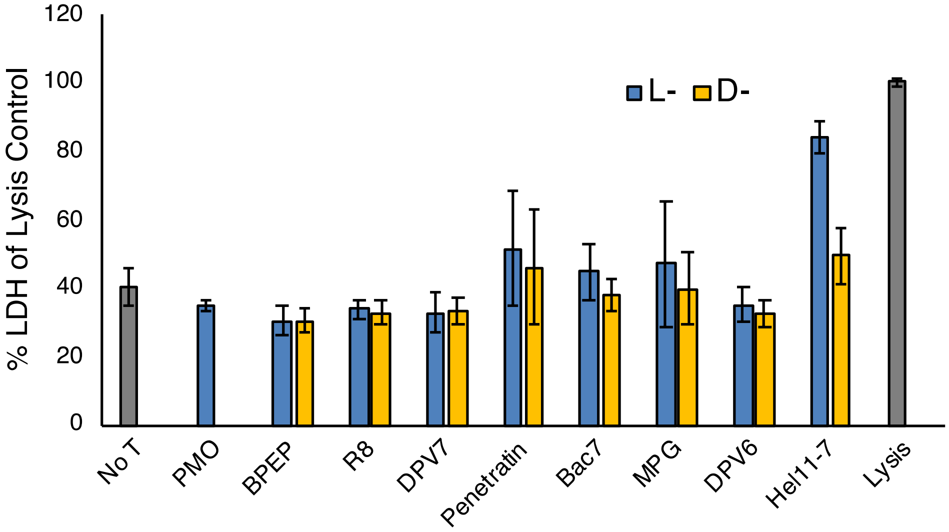

**Fig S7**. LDH Assay of first-generation peptide conjugates. Notably, PMO-Penetratin and Hel11-7 induce LDH release above background.

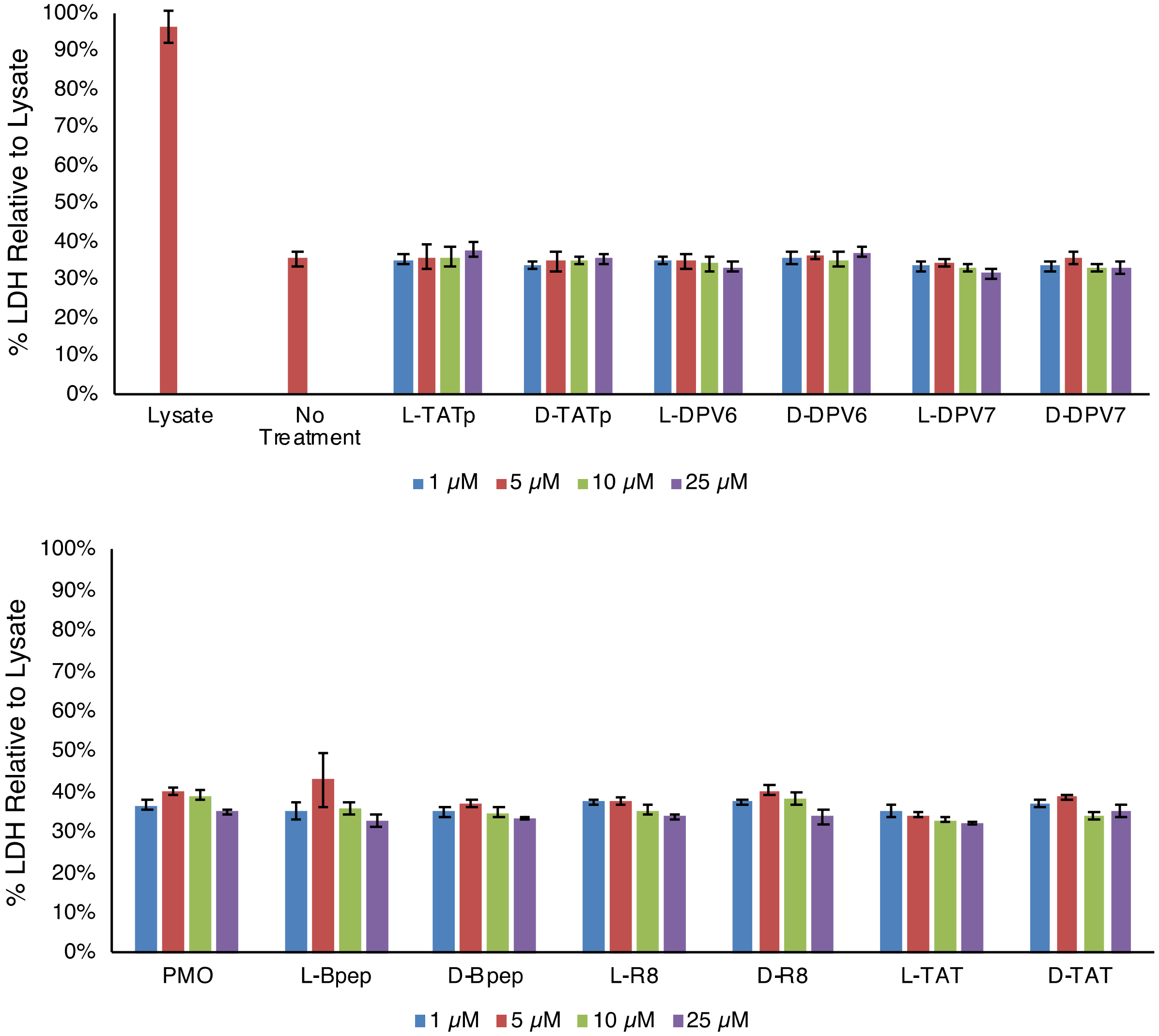

**Fig S8**. LDH release assay corresponding to EGFP assay from Fig S3.

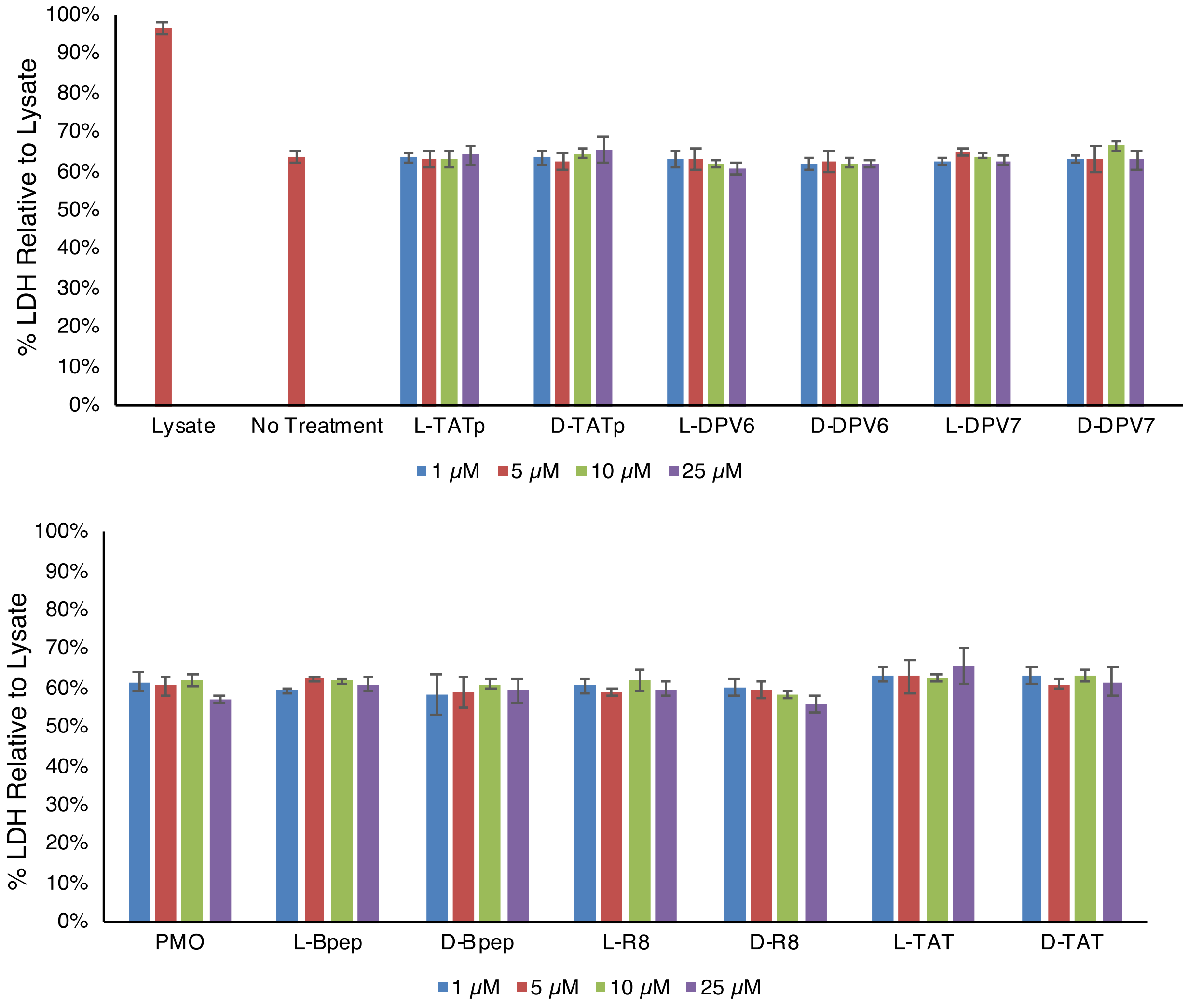

**Fig S9**. LDH release assay corresponding to EGFP assay from Fig S4.

#### 2.4 Serum Stability Assay

Each PMO-peptide was dissolved in PBS to a concentration of 1 mM, as confirmed by UV-Vis. PMO-peptide was then added to a solution of either PBS or PBS containing 25% human serum to a final concentration of 50 µM and incubated at 37 °C. 10 µL aliquots were removed at varying timepoints (t = 0, 1h, 6 h, 24 h), and quenched with 20 µL 1M guanidinium hydrochloride and 50 mM EDTA. 50 µL ice-cold acetonitrile was then added and the aliquots were flash frozen until LCMS analysis. Samples were thawed, and a portion of the aqueous layer was diluted before analysis by LC-qTOF. The mass spectrum for the PMO-peptides was analyzed by deconvolution, to best demonstrated whether the analyte had stayed intact or degraded.

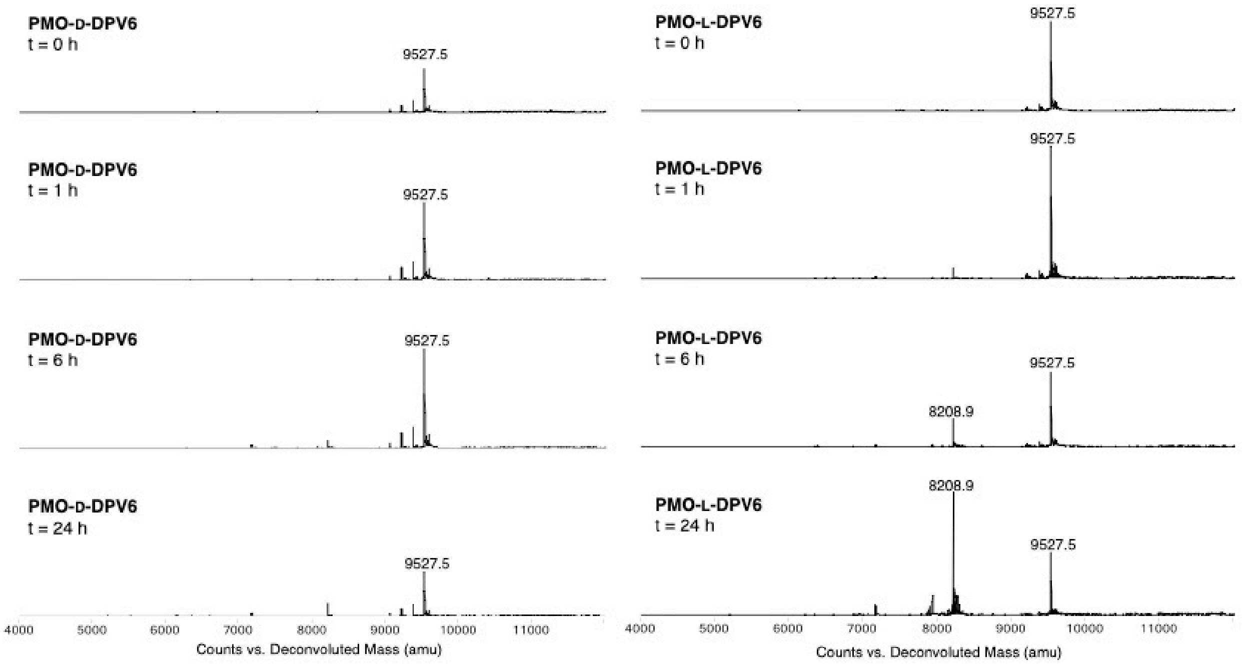

**Fig S10**. LC-MS deconvoluted mass spectrum of PMO-D or L-DPV6 following incubation in human serum.

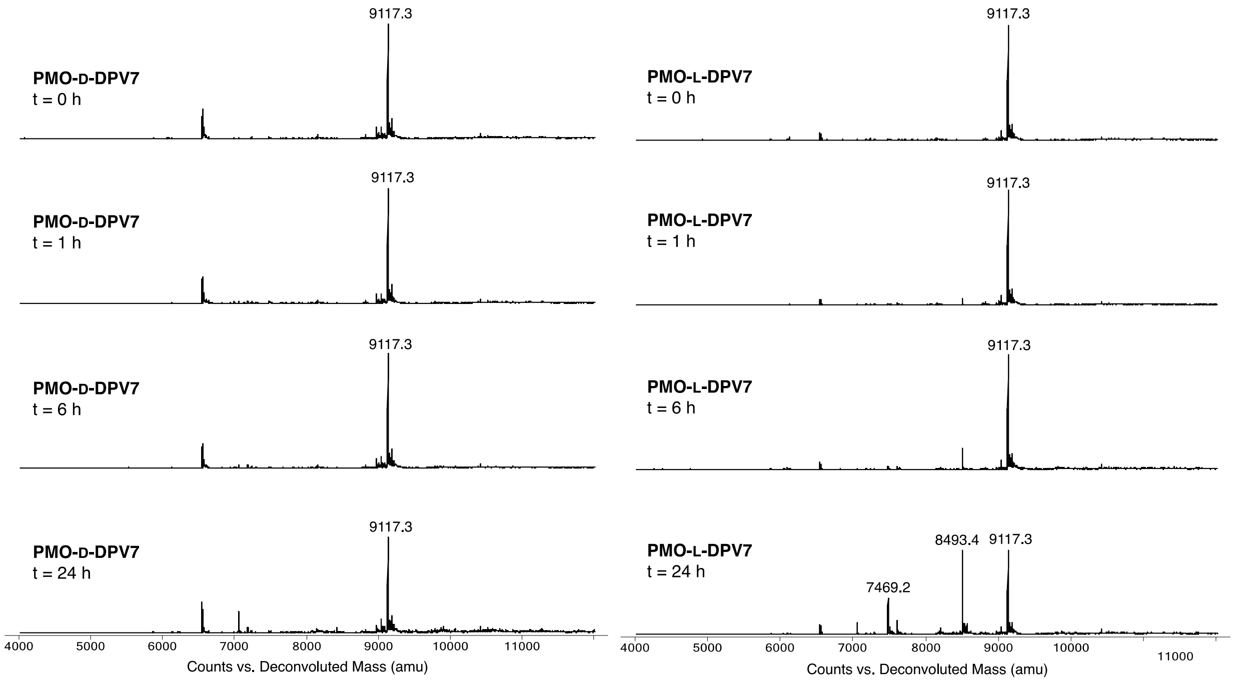

**Fig S11**. LC-MS deconvoluted mass spectrum of PMO-D or L-DPV7 following incubation in human serum.

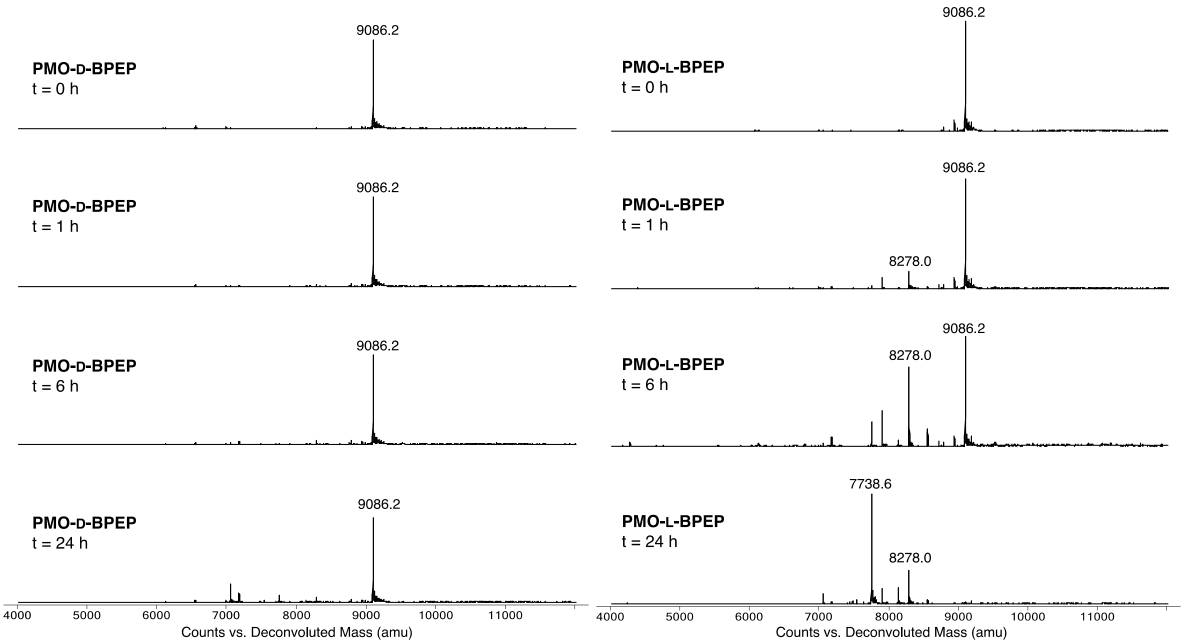

**Fig S12**. LC-MS deconvoluted mass spectrum of PMO-D or L-BPEP following incubation in human serum.

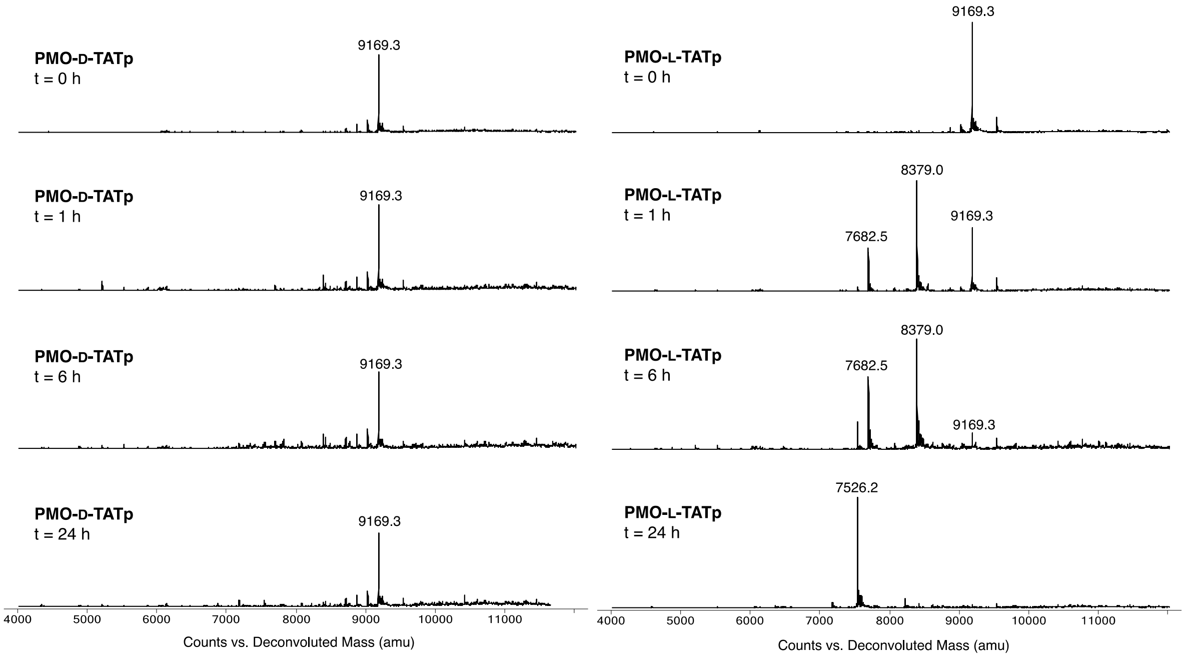

**Fig S13**. LC-MS deconvoluted mass spectrum of PMO-D or L-TATp following incubation in human serum.

#### 3. Uptake Assay

##### 3.1 General protocol

Cell treatment

Cells were plated either in 6-well or 12-well plates at a density such that they reached 80% confluency the following day. CPP or PMO-CPP stock solutions were made fresh to 1 mM in cation-free PBS, as determined by UV-Vis. Treatment solution was then prepared by adding the stock solution to cell media at the concentrations described. Two wells were left untreated as controls. The plates were then incubated at 37 °C and 5% $CO_2$ for the designated time. For the experiment to arrest energy-dependent uptake, the plate was incubated at 4 °C. Following incubation, the cells were washed three times with media, followed by 0.1 mg/mL Heparin in PBS for 5 min. Supernatant was aspirated and cells were lifted by incubating in trypsin-EDTA for 10 min at 37 °C. Trypsin was quenched by adding cell media, and cells were transferred to Eppendorf tubes and pelleted at 500 rcf for 3 min. Pellets were washed by mixing with PBS, repeated twice.

Lysis

To acquire whole cell lysate, 50 μL RIPA (1x RIPA, protease inhibitor cocktail, water) was added to the cell pellet, mixed gently, and placed on ice for 1 h. To extract the cytosol, 50 μL digitonin buffer (0.05 mg/mL digitonin, 250 mM sucrose, PBS) was added to a cell pellet, mixed very gently, and placed on ice for 10 min. Samples were then pelleted by centrifugation at 16,000 rcf for 5 min. Supernatants were transferred to new Eppendorf tubes and kept on ice. Extracted protein was quantified using Pierce Rapid Gold BCA Protein Assay Kit (Thermo Fisher). 10 μg protein from each sample was then analyzed by SDS-PAGE gel for 35 min at 165 V and then transferred to a nitrocellulose membrane soaked in 48 mM Tris, 39 mM glycine, 0.0375% SDS, 20% methanol using a TransBlot Turbo Semi-Dry Transfer Unit (BioRad) for 7 mins. The membrane was blocked at 4 °C overnight in LI-Cor Odyssey blocking buffer (PBS). The membrane was then immunostained for 1 h with anti-Erk1/2 and anti-Rab5 in TBST at room temperature. After incubation, the membrane was washed three times with TBST and incubated with the appropriate secondary antibody in TBST for 1 h at room temperature, then washed with TBST. Finally, the membrane was incubated with streptavidin-HRP for 1 h and washed with TBST. To visualize HRP, the membrane was treated with SuperSignal West Pico PLUS chemiluminescent substrate (Thermo Fisher) immediately before imaging on a ChemiDoc MP Imaging System (Bio-Rad).

MALDI-TOF

The remaining cell extracts were then used for affinity capture and MALDI-TOF analysis, following an adapted protocol.[6] 10 µL Dynabeads™ MyOne™ Streptavidin T1 (Thermo Fisher) were transferred to tubes in a magnet stand and washed with PBS. Cell extracts were added to the corresponding bead-containing tube and rotated at 4 °C overnight. One tube contained beads that were added to an equimolar solution of peptide conjugates used in the experiment. To insure the same equivalency are in the control tube as used in the experiment, this control solution was taken directly from the combined stock solution used in the initial cell treatment. The following day, the beads slurries were washed with a series of buffers: 2 x 100 µL Buffer A (50 mM Tris-HCl (pH 7.4) and 0.1 mg/mL BSA), 2 x 100 µL Buffer B (50 mM Tris-HCl (pH 7.4), 0.1 mg/mL BSA, and 0.1% SDS), 2 x 100 µL Buffer C (50 mM Tris-HCl (pH 7.4), 0.1 mg/mL BSA, and 1 M NaCl), and 2 x 100 µL water. Beads were then incubated with 100 µL of 1 mM biotin for 2 min, before washed with 5 x 50 µL water. Supernatant was removed and the beads were brought up in 3 µL MALDI matrix (saturated alpha-Cyano-4-hydroxycinnamic acid, CHCA) and transferred to the MALDI plate to dry. Beads were analyzed by MALDI-ToF on a high-resolution Bruker Autoflex LRF Speed mass spectrometer in linear positive mode.

Relative concentrations of peptides in the mixture were determined as follows. Analytes in a mixture ionize according to their response factor (F). F was determined by normalizing the intensities of each analyte to one analyte in the control sample, where the concentration of each analyte is arbitrarily set to 1. The values of F is then used in the experimental spectra containing the same mixture of analytes to determine their relative concentrations.

$$\frac{I_x}{[X]} = F\left(\frac{I_s}{[S]}\right)$$

#### 3.2 Supplementary Uptake Data

We confirmed by orthogonal means that the PMO-peptide conjugates entered via energy-dependent uptake and that outer membrane-bound conjugates do not contaminate lysate samples. First, an EGFP assay determined that for both PMO-D- and L-DPV7, incubation at reduced temperature negatively impacted PMO delivery (Fig S14). Then, HeLa cells were incubated with three PMO-D-CPPs at 37 °C or 4 °C before washing and lysis as before. Analysis by Western blot shows presence of both cytosolic and endosomal markers in both whole cell lysates, but shows a marked absence of biotinylated construct in the 4C condition by Streptavidin labeling. Analysis of these samples by MALDI-TOF also shows significantly reduced signal in the 4 °C condition compared to 37C, where only PMO-D-DPV7 is detected at reduced temperature. At the same time, no construct was detected in the 4 °C cytosolic condition. By using the constructs response factor (F) from the equimolar condition, we found that PMO-D-DPV7 had the highest relative intracellular concentration, although the three constructs had very similar concentrations.

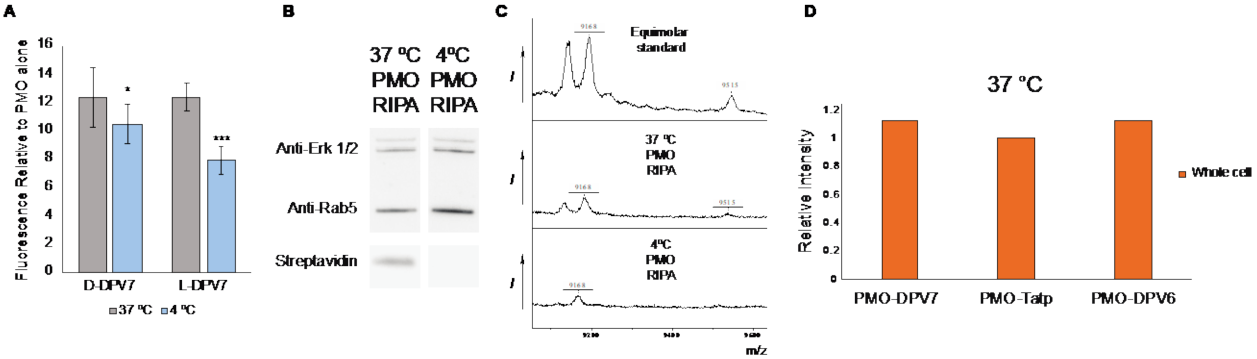

**Fig S14**. **Incubation at 4 °C inhibits internalization of PMO-D-CPPs.** (A) Bar graph showing fluorescence relative to PMO alone as measured in the EGFP activity assay, comparing uptake of D- and L-DPV7 at 5 µM at 37 °C to 4 °C. Bars represent group mean ± SD, N = 3 distinct samples from a single biological replicate (*p<0.05, ***p<0.0005) (B) Western blot indicating the presence of PMO-biotin-CPPs in whole cell lysate following 1 h incubation at 37 °C, but not at 4 °C. (C) MALDI-TOF mass spectra corresponding to equimolar standard (top), whole cell lysate following treatment at 37 °C (middle) and 4 °C (bottom). Only PMO-D-DPV7 is observable after incubation at 4 °C, indicating that uptake of these conjugates is arrested at low temperature. (D) Graph showing relative concentration of the three PMO-D-CPPs in whole cell lysate following incubation at 37 °C for 1 h, performed in singlicate.

Interestingly, reduced temperature did not inhibit uptake of the biotinylated-CPPs without the oligonucleotide cargo, but rather equalized the relative concentration of the whole cell and cytosolic fractions. In the same manner as the experiment with PMO-D-CPPs, biotin-D-DPV7,

TATp, and DPV6 were incubated in HeLa cells at 37C and 4C. Cytosol and whole cell lysate were extracted, confirmed by Western blot (SI). Samples were then analyzed by MALDI and the relative concentrations were determined, normalized to TATp. At 37C, the determined relative calculations vary between the whole cell and cytosolic samples; DPV7 appears to have the highest concentration in the cytosol (Fig S15). However, at reduced temperature, the relative concentrations are nearly identical between the lysates. This observation is unsurprising when its considered that low temperature arrests endocytosis, meaning that the only material inside the cell likely entered through passive diffusion through the membrane to directly access the cytosol.[7,8] It is also unsurprising that CPPs with a single small biotin label are able to directly translocate, whereas peptides attached to macromolecules such as PMO cannot.[9] These experiments demonstrate that this method, in addition to determining relative internal concentration, is useful for investigating mechanisms of uptake.

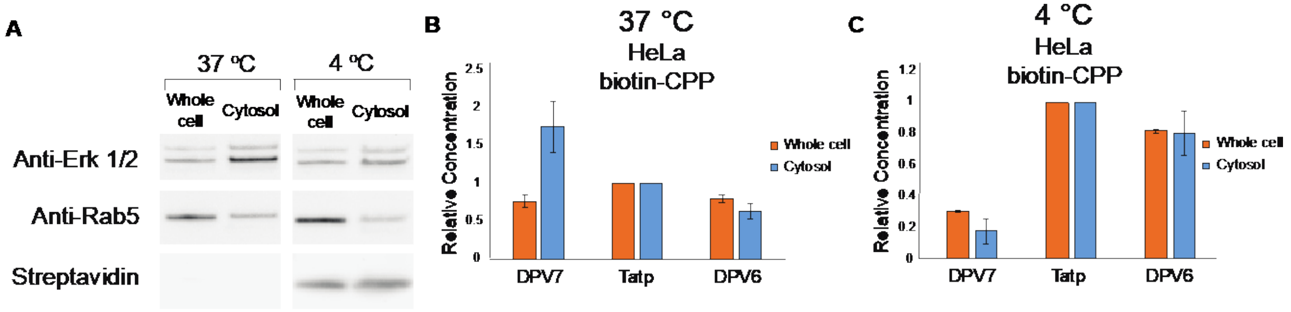

**Fig S15. Biotin-D-CPPs were incubated with HeLa cells at 37 °C or 4 °C for 4 h.** (A) Western blot showing cytosolic extraction of the samples analyzed by MALDI. Concentrations of biotin-peptide relative to TATp in the whole cell and cytosolic extracts following 4 h treatment at (B) 37 °C and at (C) 4 °C. Bars show group mean ± SD, N = 2 distinct samples from a single biological replicate. Experiment was repeated with shorter incubation time with similar results, shown in Fig S16.

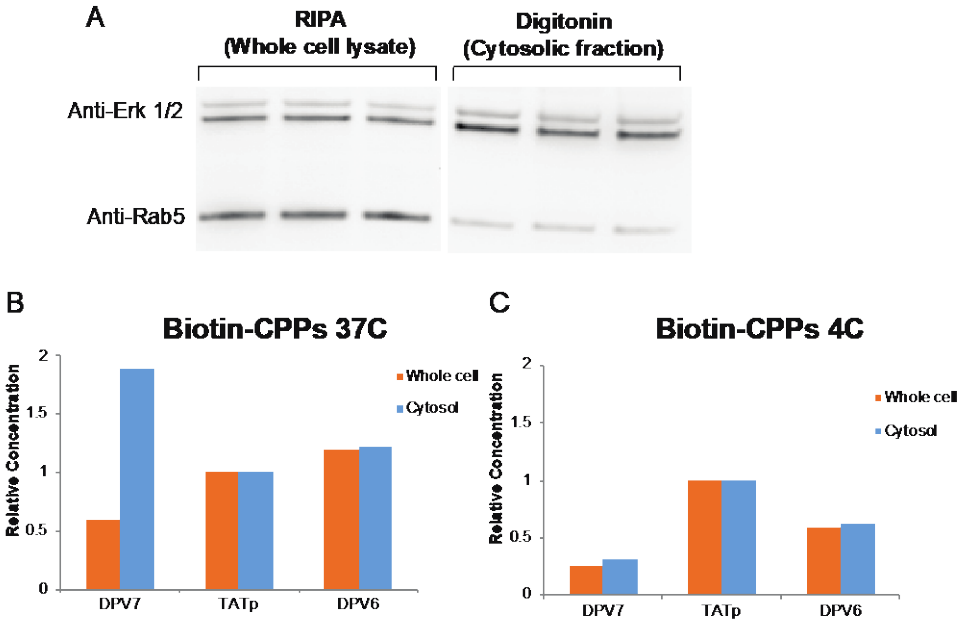

**Fig S16. Additional Temperature experiment**. Biotin-D-CPPs were incubated with HeLa cells at 37C or 4C for 1 h. (A) Western blot showing cytosolic extraction of the samples analyzed by MALDI. (B-C) Graph showing relative concentration of biotin-CPPs in the whole cell and cytosolic fractions following incubation at (B) 37C and (C) 4C as determined by MALDI-TOF. Bars represent biological singlicate.

#### Appendix I: Gel Images

Full gel image corresponding to that shown in Figure 4.

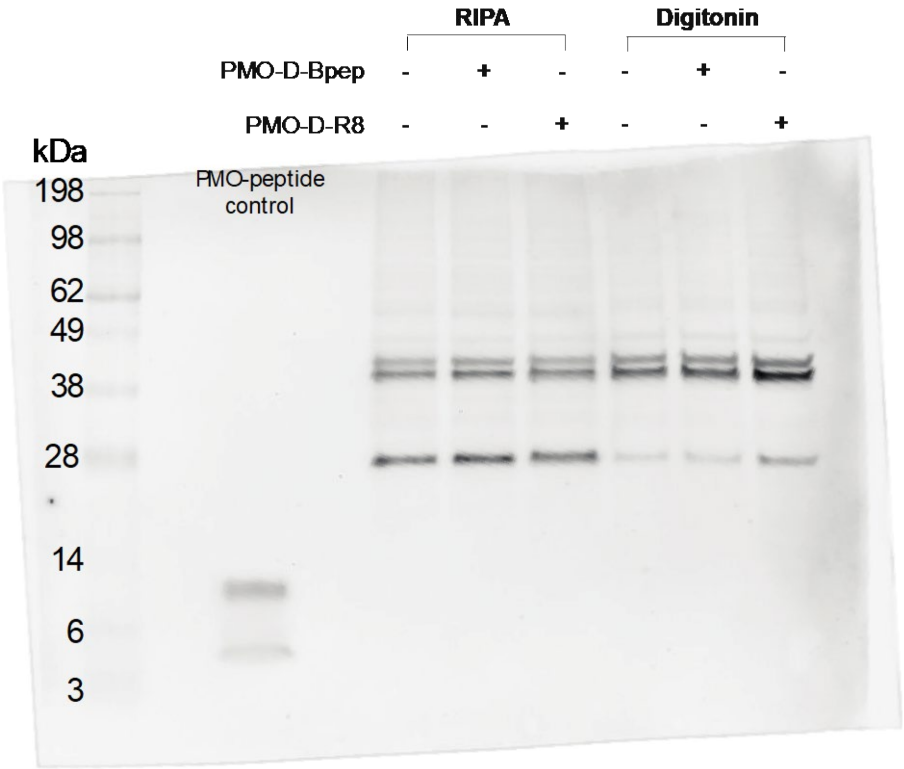

Full gel image corresponding to the experiment described in Figure 5.

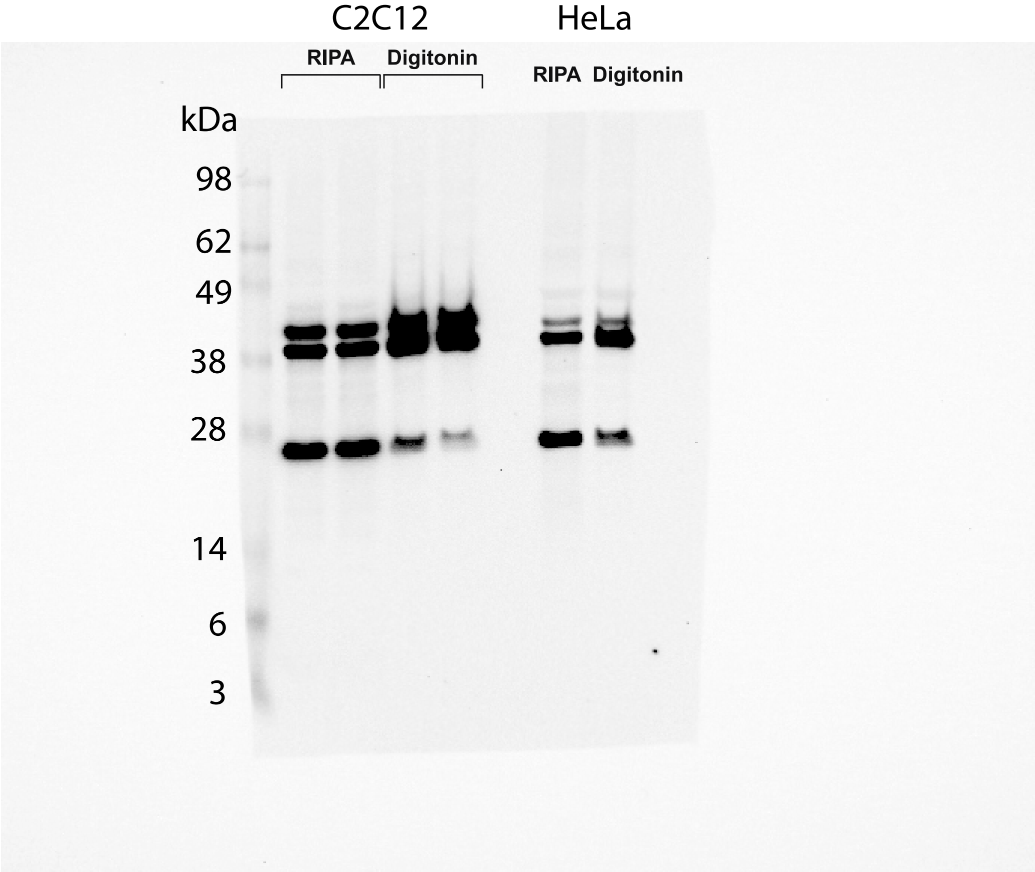

Full gel image corresponding to that shown in Figure S14 and S15. IR800 (left) and Chemiluminescence (right)

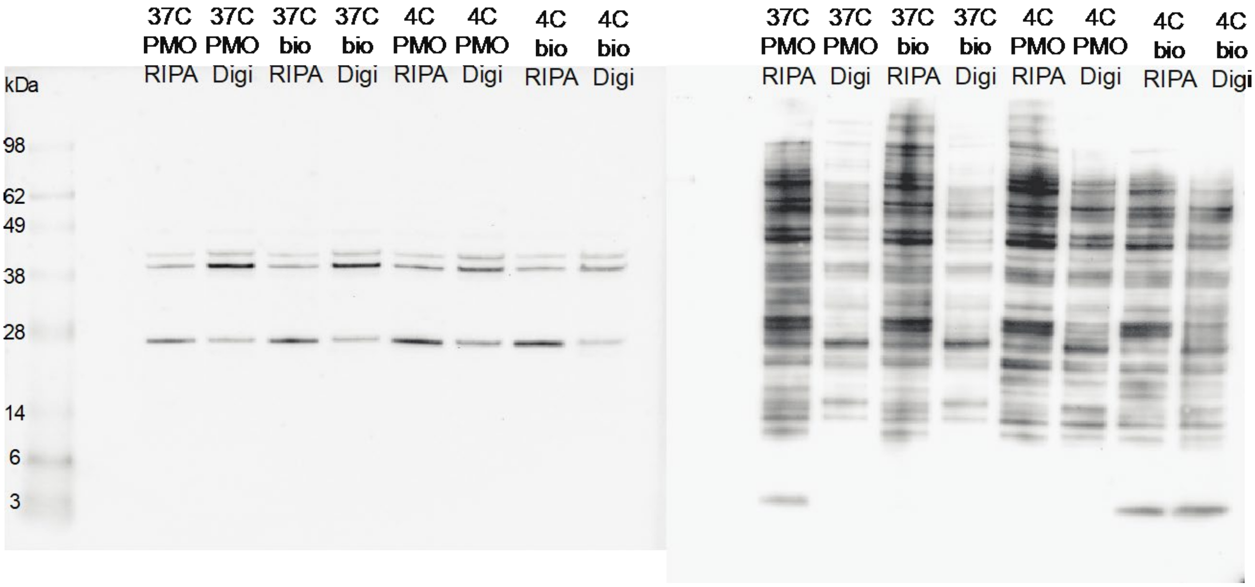

Full gel image corresponding to that shown in Figure S16

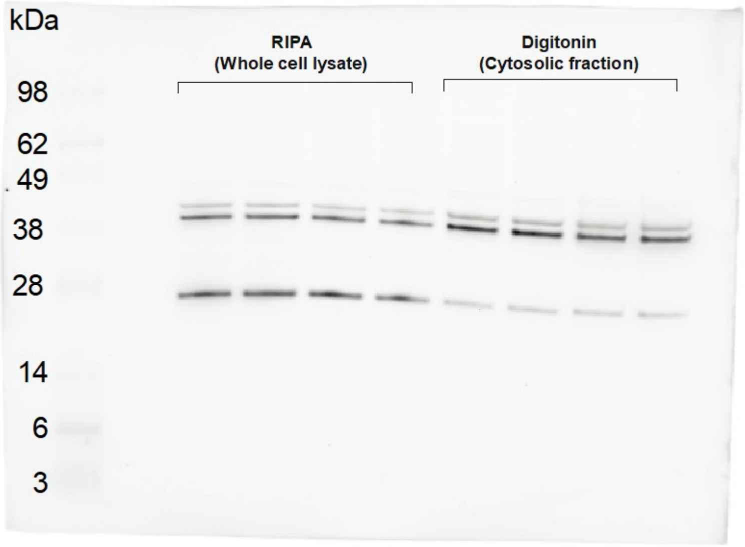

#### Appendix II: LC-MS Characterization

Peptide: Bio-D-TAT
Sequence: Biotin-Lys($N_3$)-GGKGGWRKKRRQRRR
Calculated monoisotopic mass: 2260.3 Da
Observed monoisotopic mass: 2260.4 Da

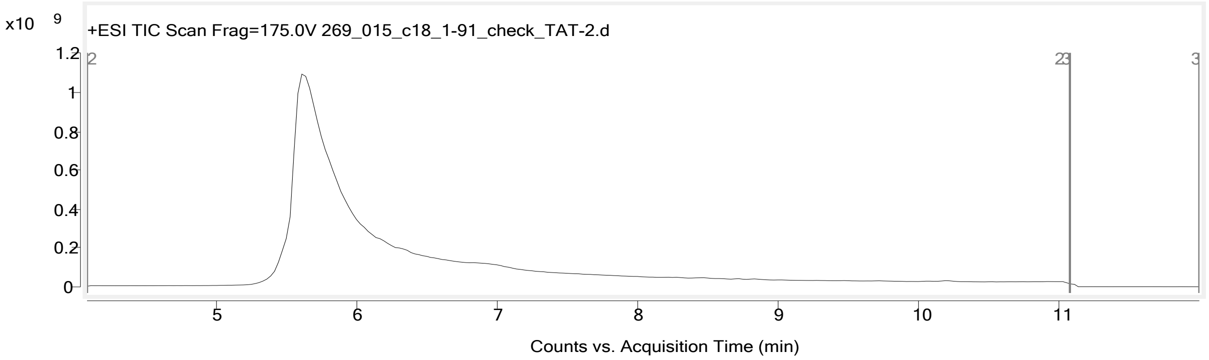

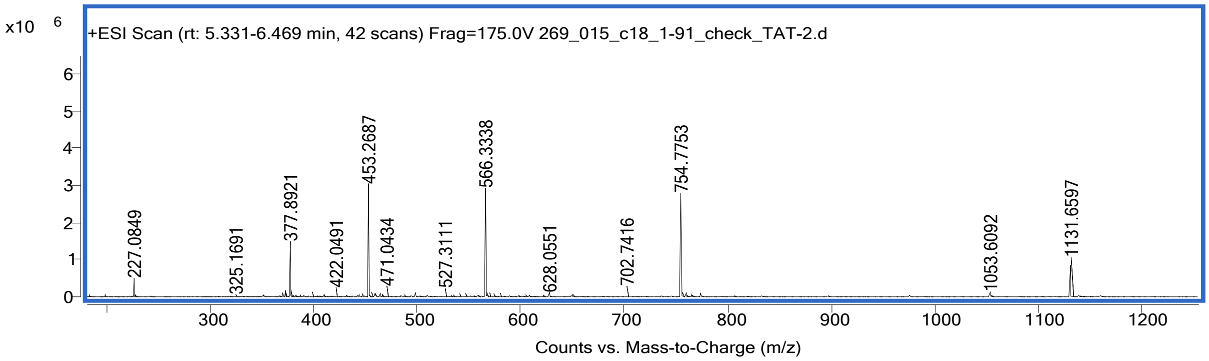

Peptide: Bio-L-TAT
Sequence: Biotin-Lys($N_3$)-GGKGGWRKKRRQRRR
Calculated monoisotopic mass: 2260.3 Da
Observed monoisotopic mass: 2260.4 Da

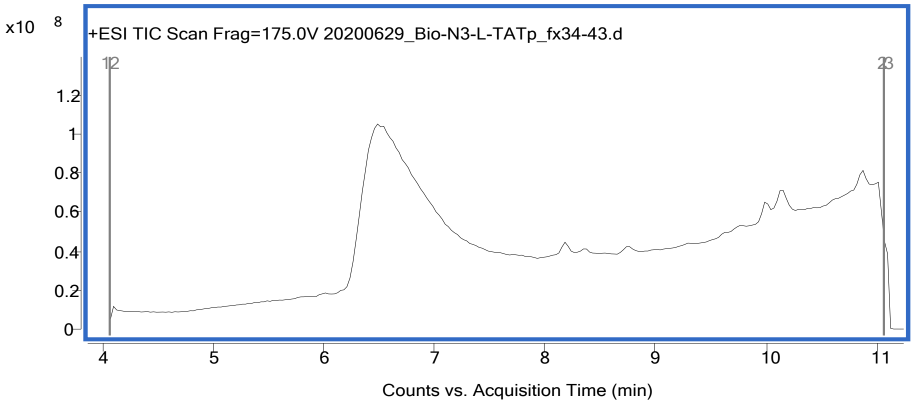

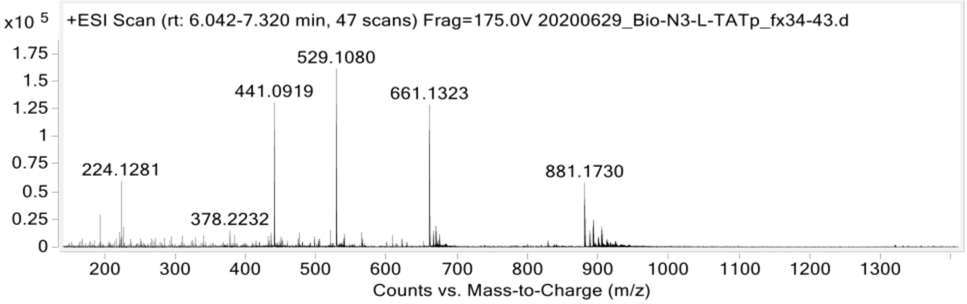

Peptide: Bio-D-TATp
Sequence: Biotin-Lys($N_3$)-GGKGGWGRKKRRQRRRPPQ
Calculated monoisotopic mass: 2639.5 Da
Observed monoisotopic mass: 2639.5 Da

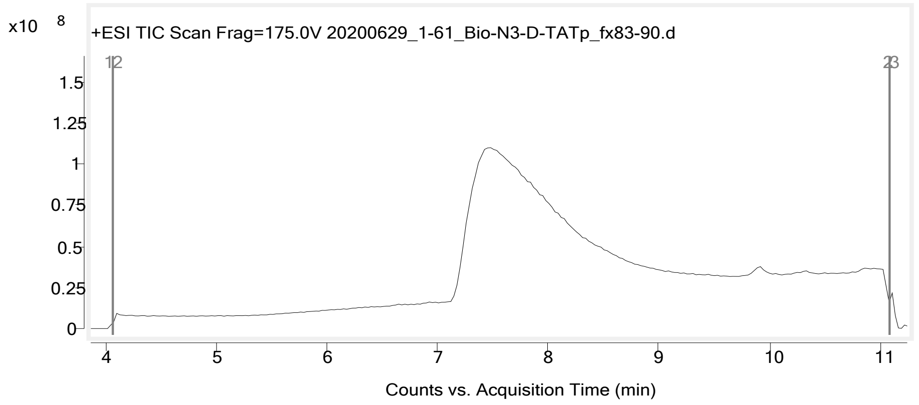

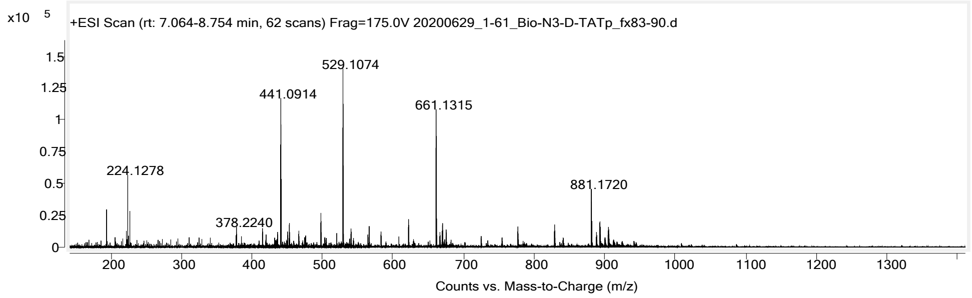

Peptide: Bio-L-TATp
Sequence: Biotin-Lys($N_3$)-GGKGGWGRKKRRQRRRPPQ
Calculated monoisotopic mass: 2639.5 Da
Observed monoisotopic mass: 2639.5 Da

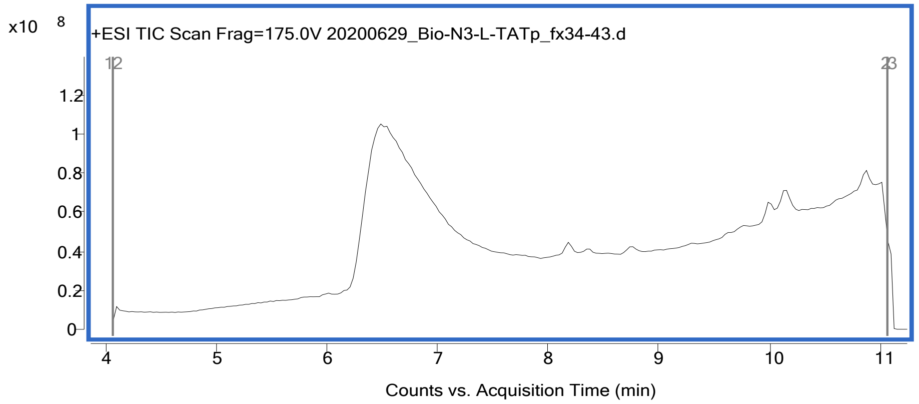

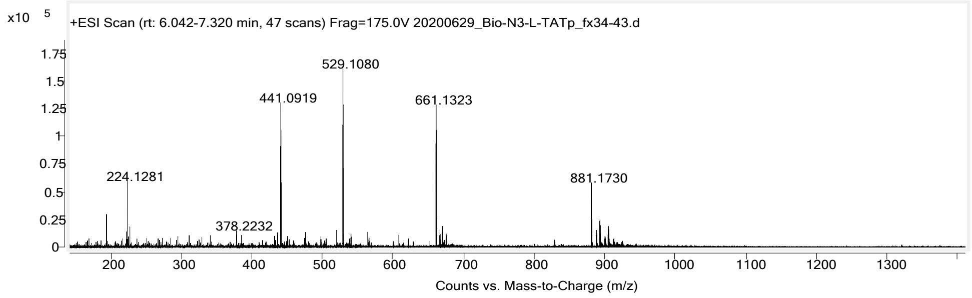

Peptide: Bio-D-DPV6
Sequence: Biotin-Lys($N_3$)-GGKGGWGRPRESGKKRKRKRLKP
Calculated monoisotopic mass: 2997.7 Da
Observed monoisotopic mass: 2997.8 Da

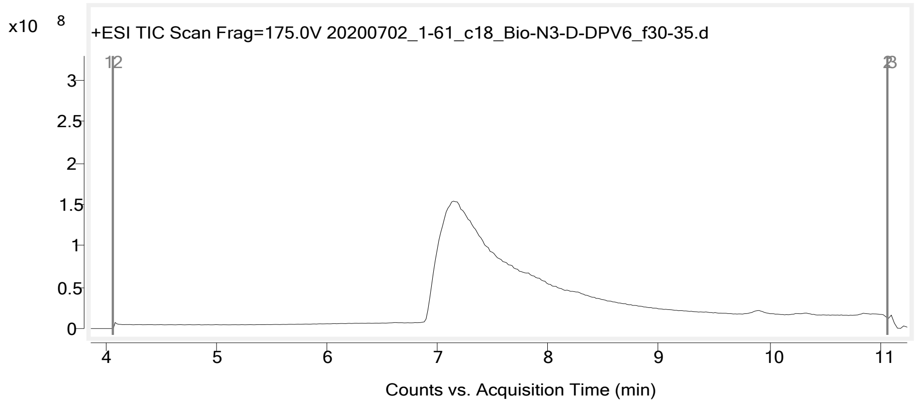

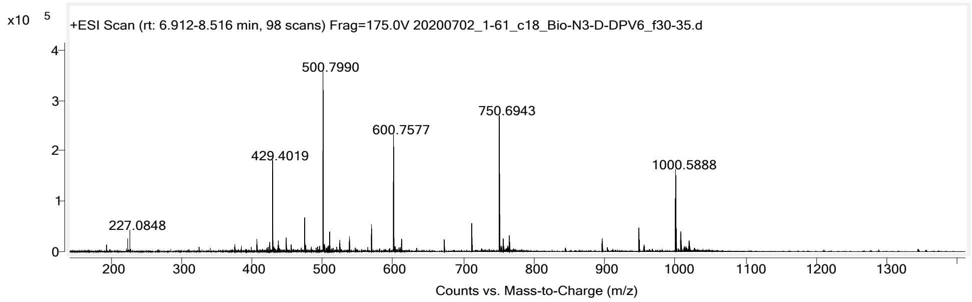

Peptide: Bio-L-DPV6
Sequence: Biotin-Lys($N_3$)-GGKGGWGRPRESGKKRKRKRLKP
Calculated monoisotopic mass: 2997.7 Da
Observed monoisotopic mass: 2997.8 Da

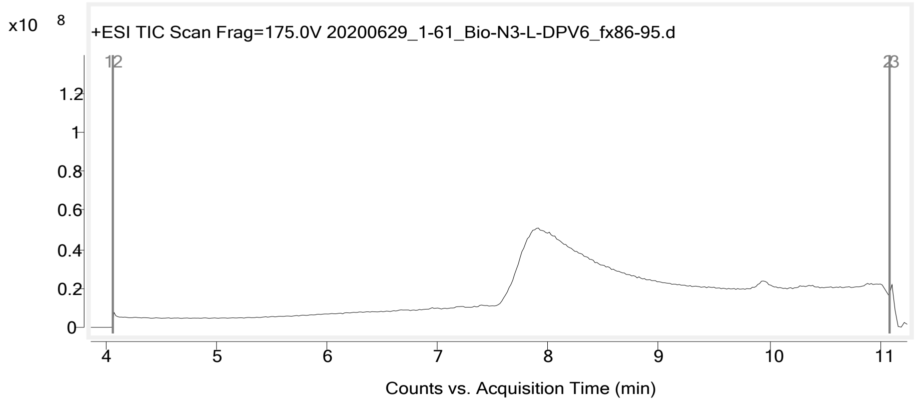

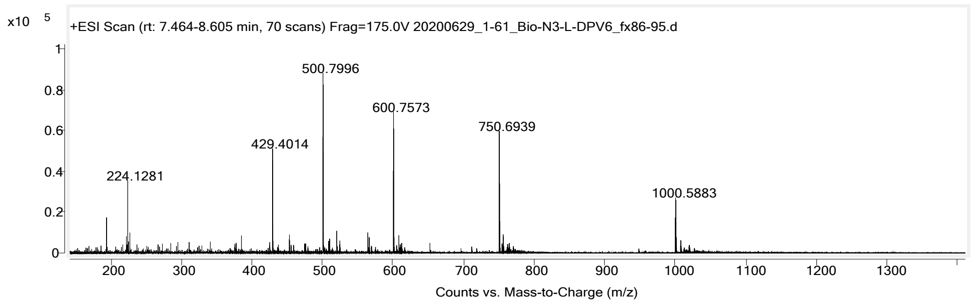

Peptide: Bio-D-DPV7
Sequence: Biotin-Lys($N_3$)-GGKGGWKRKKKGKLGKKRDP
Calculated monoisotopic mass: 2587.5 Da
Observed monoisotopic mass: 2587.5 Da

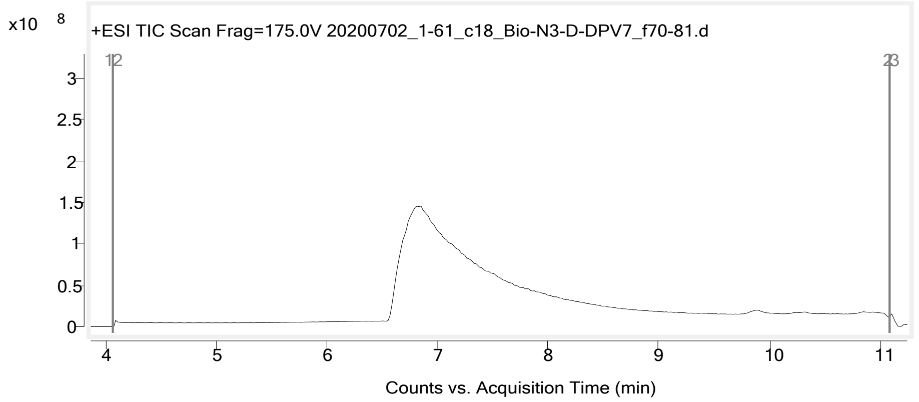

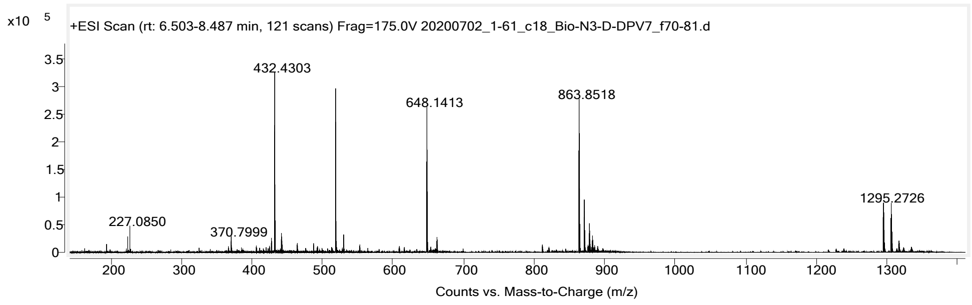

Peptide: Bio-L-DPV7
Sequence: Biotin-Lys($N_3$)-GGKGGWKRKKKGKLGKKRDP
Calculated monoisotopic mass: 2587.5 Da
Observed monoisotopic mass: 2587.5 Da

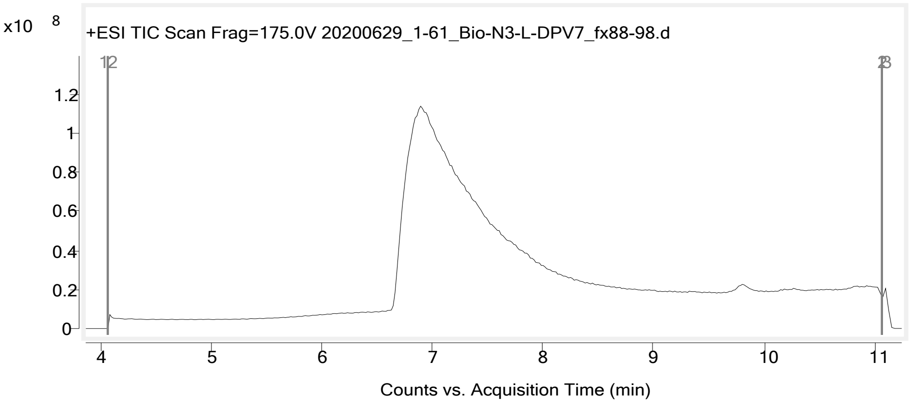

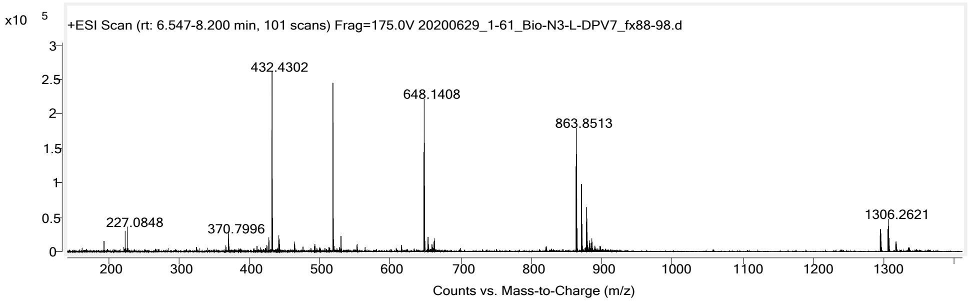

Peptide: Bio-D-BPEP
Sequence: Biotin-Lys($N_3$)-GGKGGWRXRRBRRXRRBR
Calculated monoisotopic mass: 2556.5 Da
Observed monoisotopic mass: 2556.5 Da

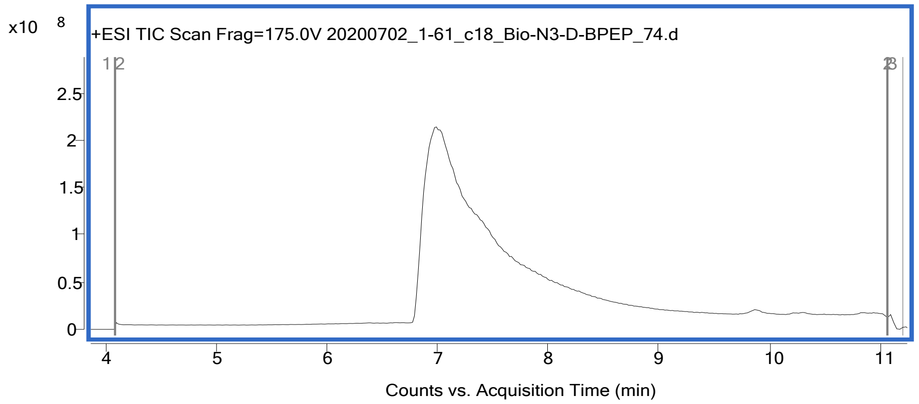

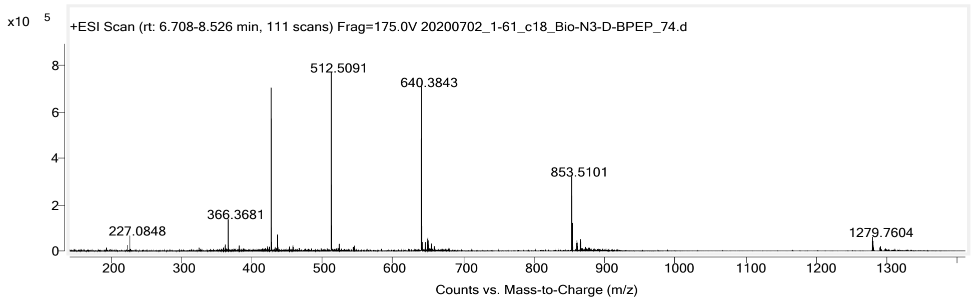

Peptide: Bio-L-BPEP
Sequence: Biotin-Lys($N_3$)-GGKGGWRXRRBRRXRRBR
Calculated monoisotopic mass: 2556.5 Da
Observed monoisotopic mass: 2556.5 Da

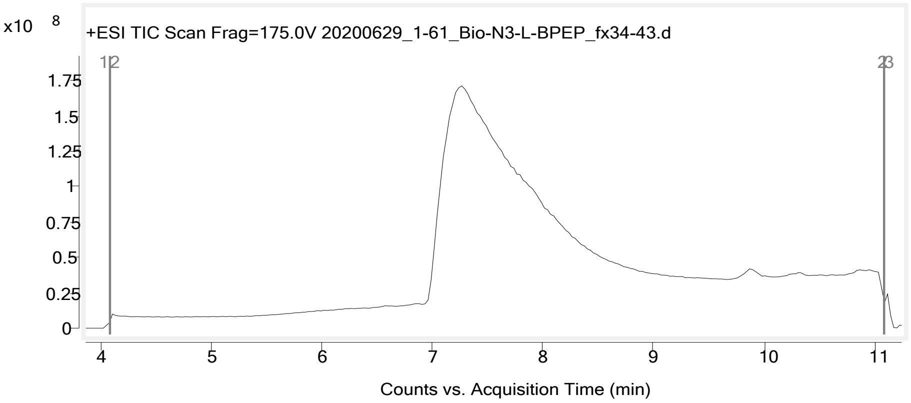

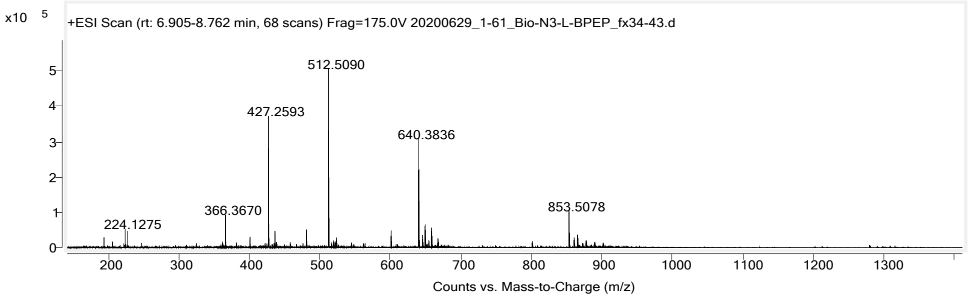

Peptide: Bio-D-R8
Sequence: Biotin-Lys($N_3$)-GGKGGWRRRRRRRR
Calculated monoisotopic mass: 2188.3 Da
Observed monoisotopic mass: 2188.3 Da

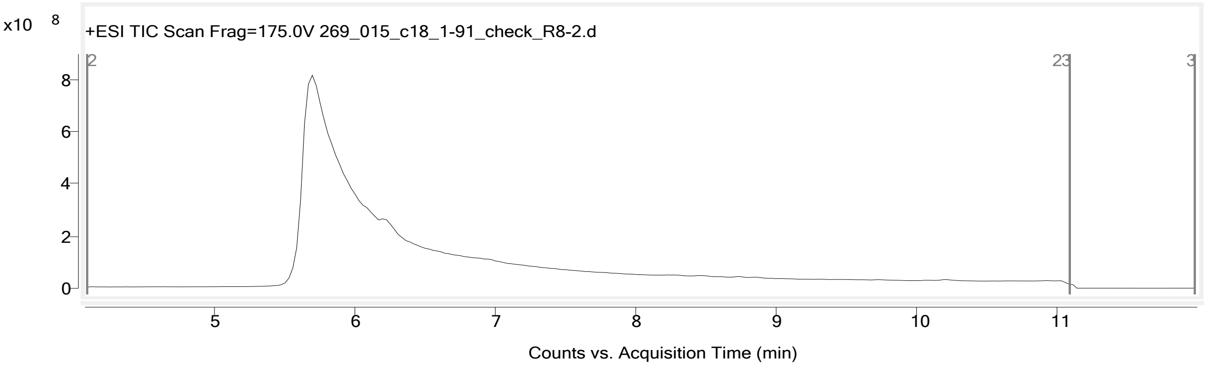

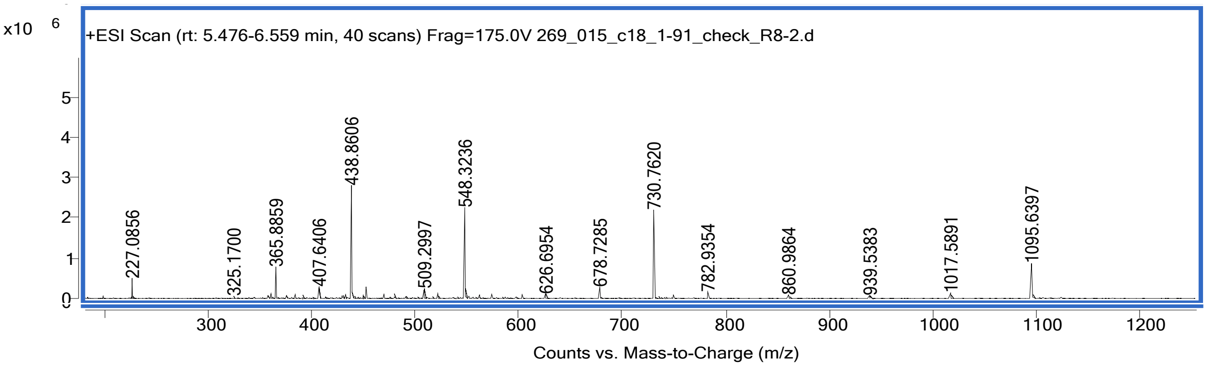

Peptide: Bio-L-R8
Sequence: Biotin-Lys($N_3$)-GGKGGWRRRRRRRR
Calculated monoisotopic mass: 2188.3 Da
Observed monoisotopic mass: 2188.3 Da

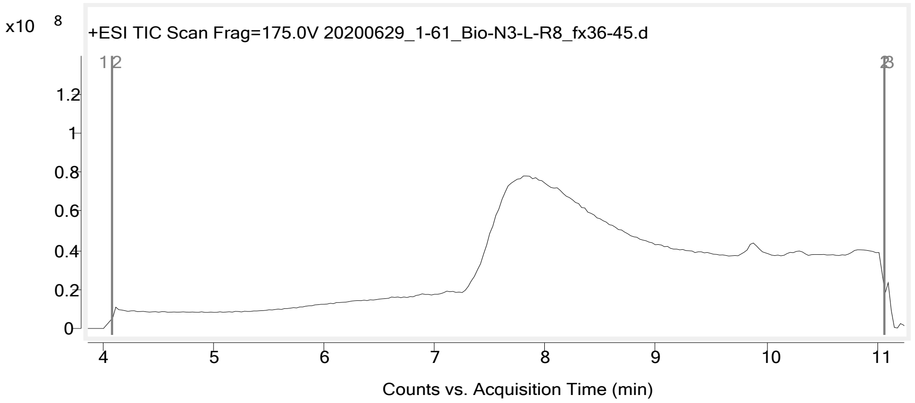

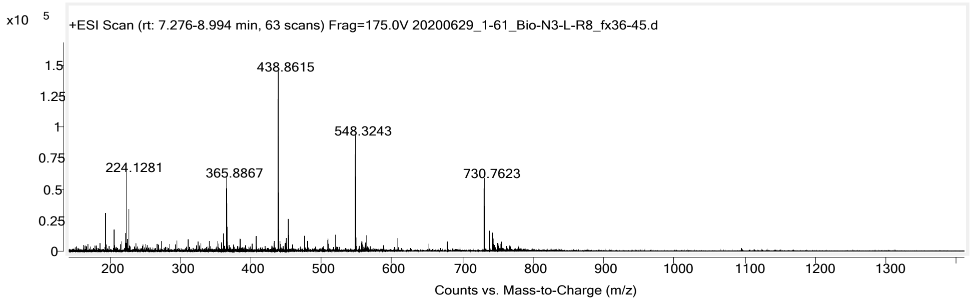

Peptide: PMO-Bio-D-TAT
Sequence: Biotin-Lys(PMO)-GGKGGWRKKRRQRRR
Calculated mass:8789.6 Da
Observed mass: 8789.8 Da

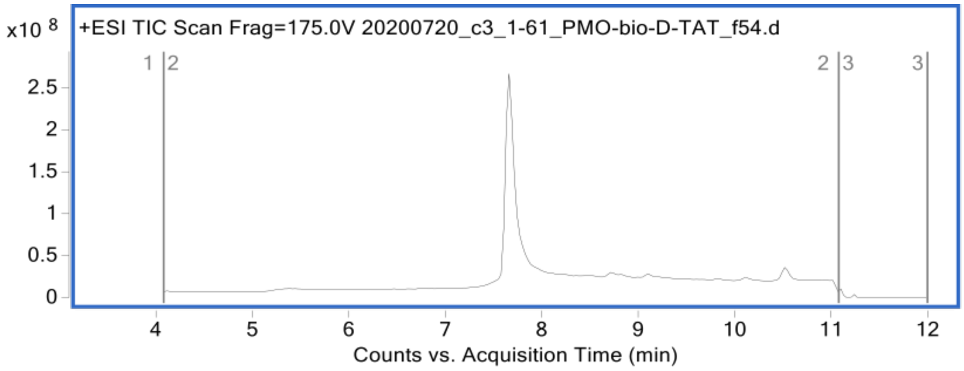

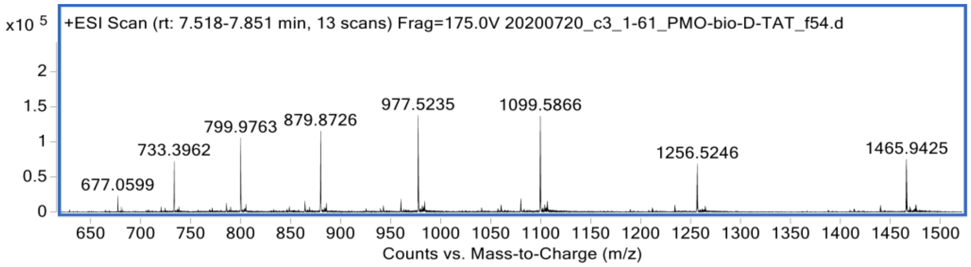

Peptide: PMO-Bio-L-TAT
Sequence: Biotin-Lys(PMO)-GGKGGWRKKRRQRRR
Calculated mass: 8789.6 Da
Observed mass: 8789.7 Da

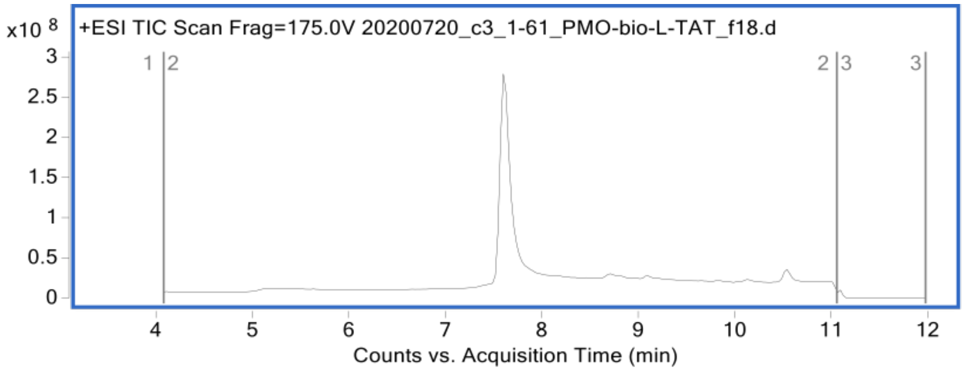

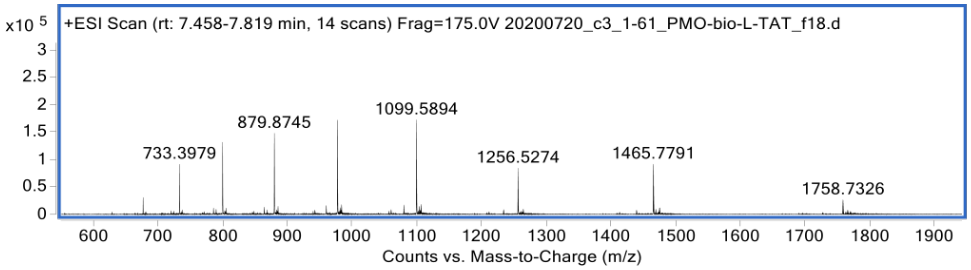

Peptide: PMO-Bio-D-TATp
Sequence: Biotin-Lys(PMO)-GGKGGWGRKKRRQRRRPPQ
Calculated mass: 9169.0 Da
Observed mass: 9169.1 Da

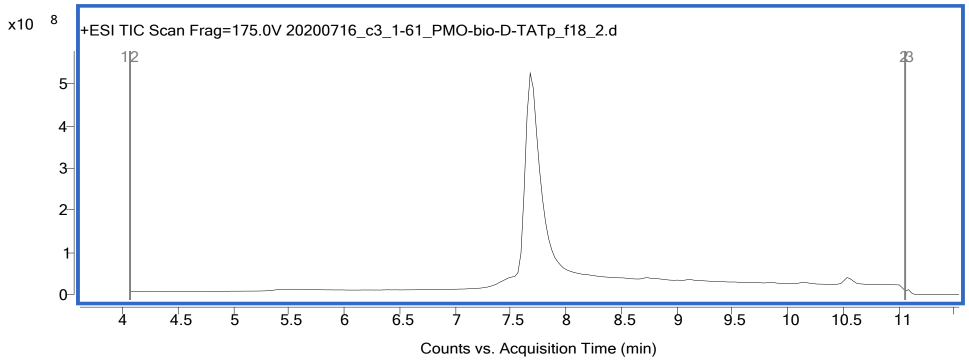

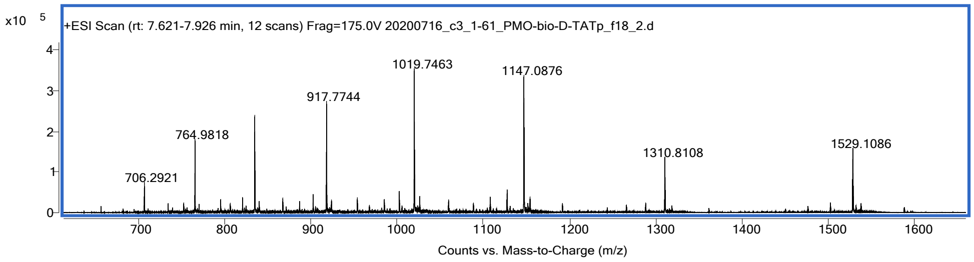

Peptide: PMO-Bio-L-TATp
Sequence: Biotin-Lys(PMO)-GGKGGWGRKKRRQRRRPPQ
Calculated mass: 9169.0 Da
Observed mass: 9169.1 Da

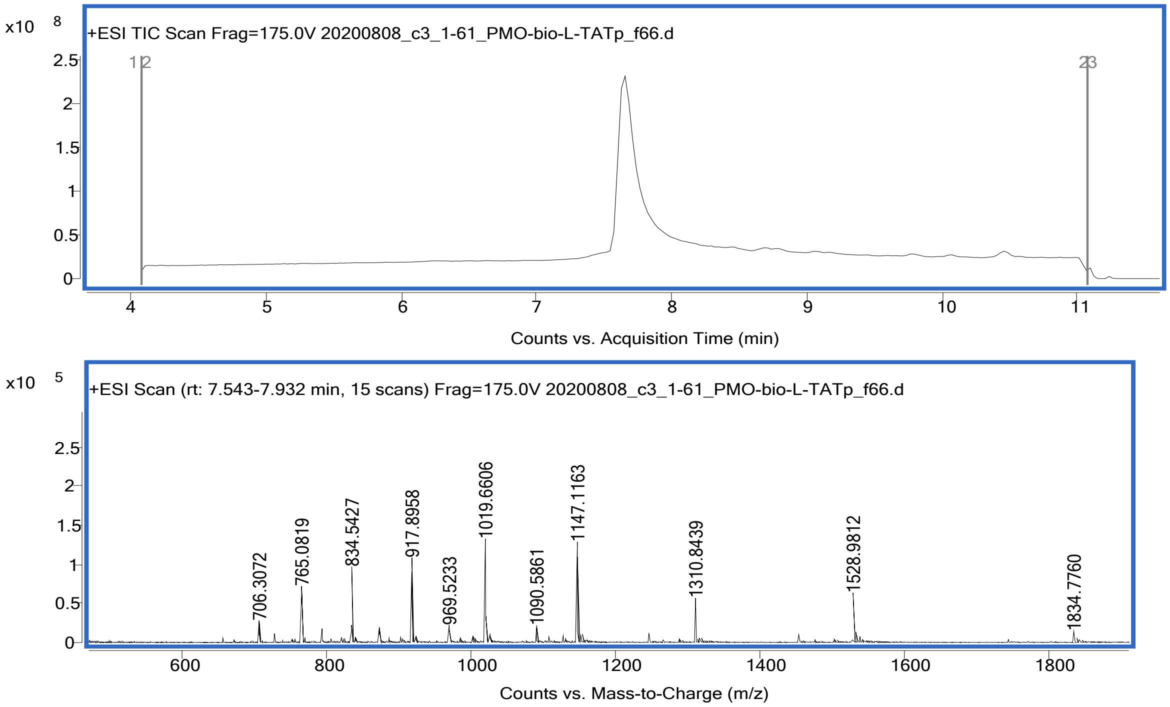

Peptide: PMO-Bio-D-DPV6
Sequence: Biotin-Lys(PMO)-GGKGGWGRPRESGKKRKRKRLKP
Calculated mass: 9527.5 Da
Observed mass: 9527.2 Da

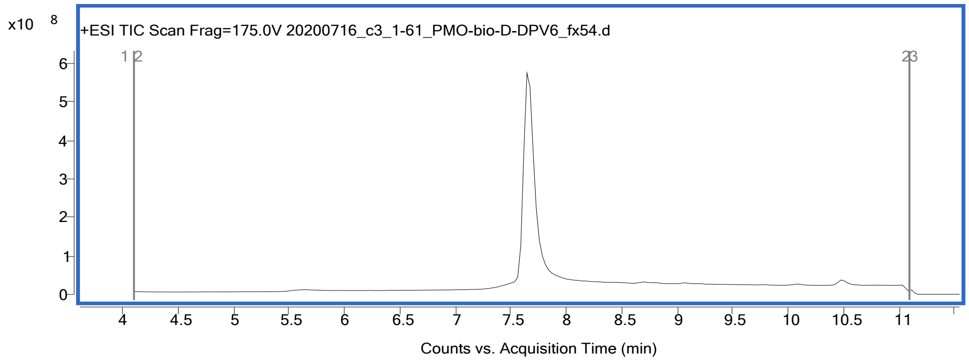

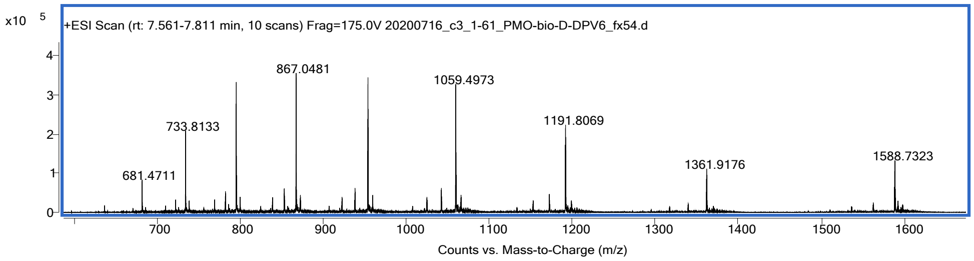

Peptide: PMO-Bio-L-DPV6
Sequence: Biotin-Lys(PMO)-GGKGGWGRPRESGKKRKRKRLKP
Calculated mass: 9527.5 Da
Observed mass: 9527.4 Da

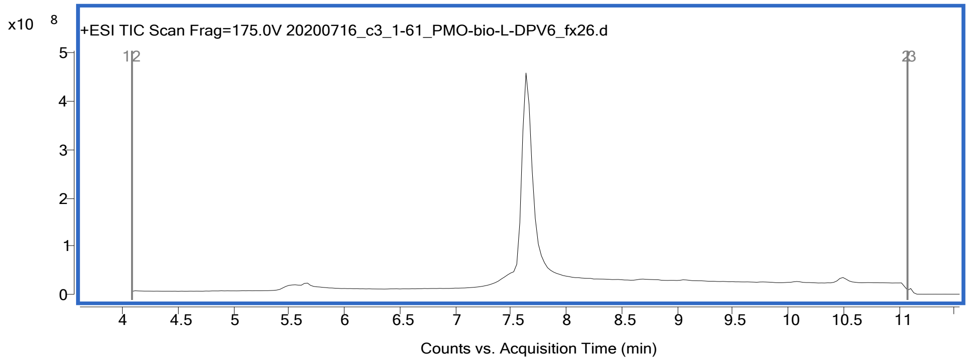

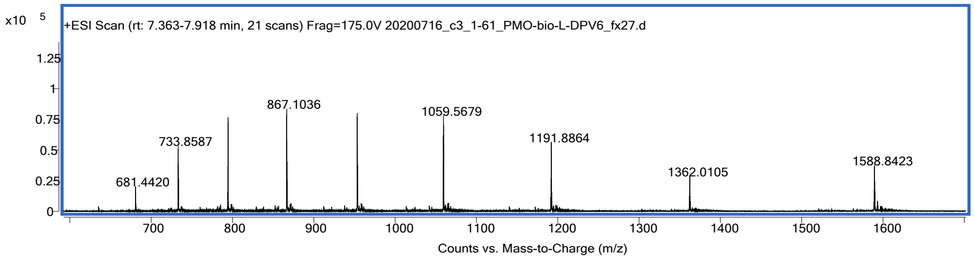

Peptide: PMO-Bio-D-DPV7
Sequence: Biotin-Lys(PMO)-GGKGGWKRKKKGKLGKKRDP
Calculated mass: 9117.1 Da
Observed mass: 9117.1 Da

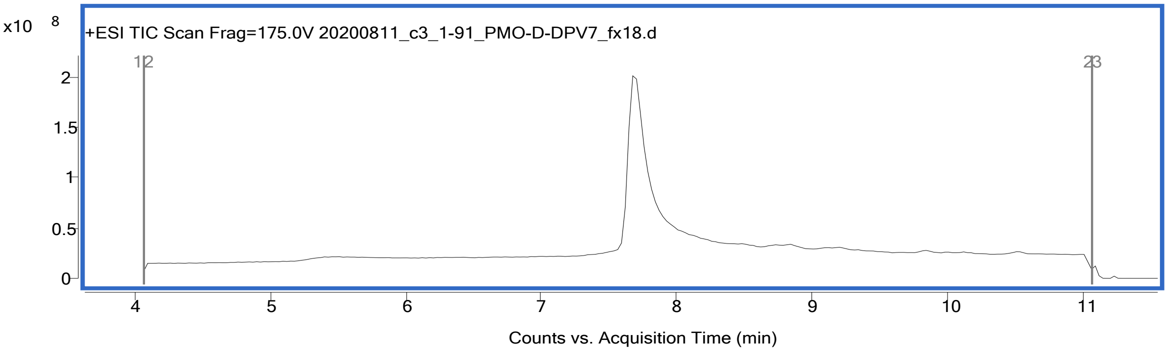

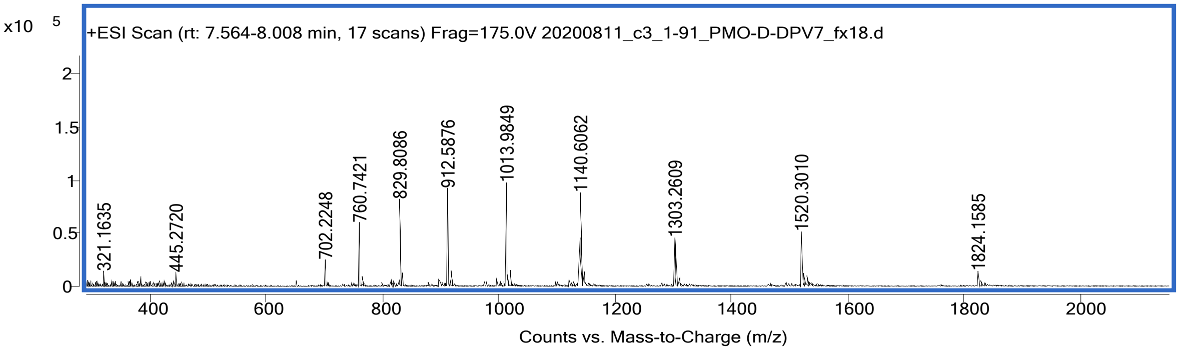

Peptide: PMO-Bio-L-DPV7
Sequence: Biotin-Lys(PMO)-GGKGGWKRKKKGKLGKKRDP
Calculated mass: 9117.1 Da
Observed mass: 9117.1 Da

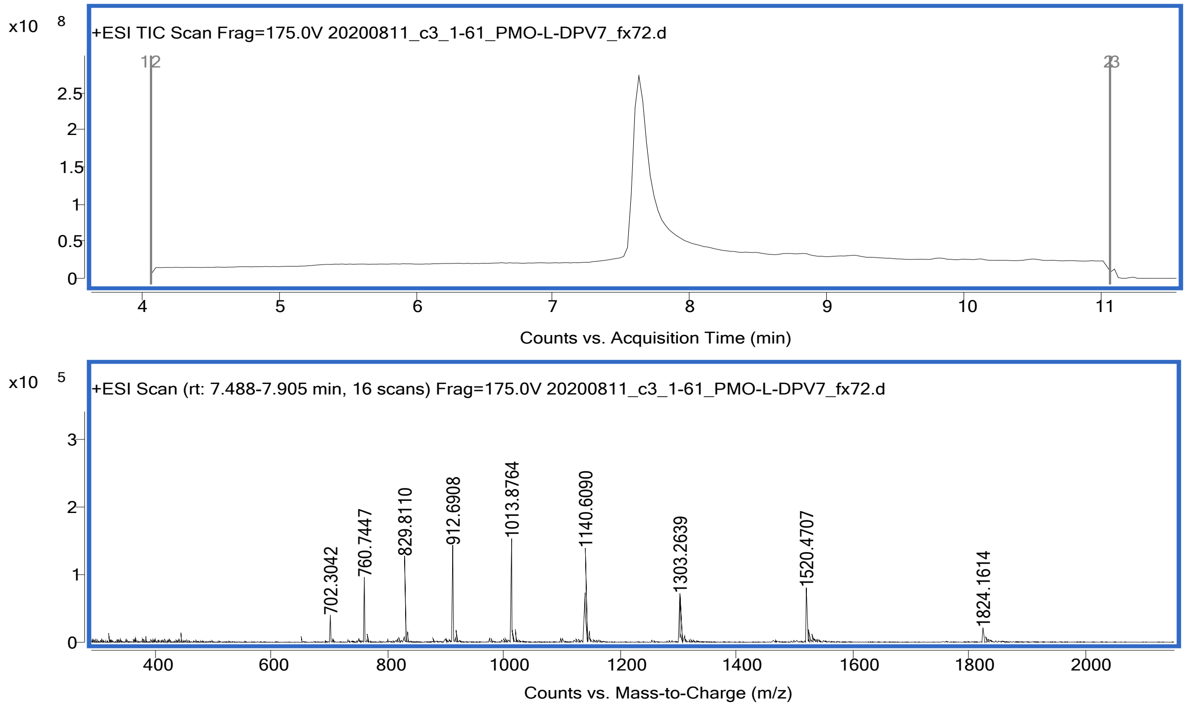

Peptide: PMO-Bio-D-BPEP
Sequence: Biotin-Lys(PMO)-GGKGGWRXRRBRRXRRBR
Calculated mass: 9086.0 Da
Observed mass: 9086.1 Da

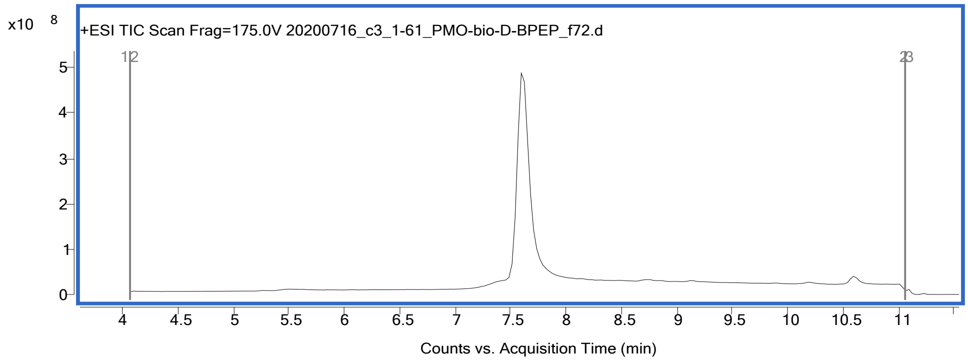

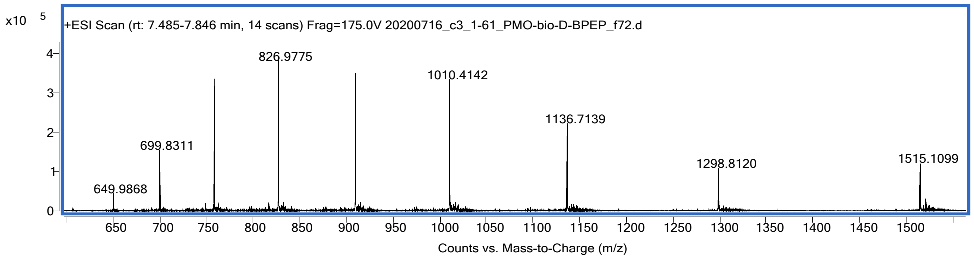

Peptide: PMO-Bio-L-BPEP
Sequence: Biotin-Lys(PMO)-GGKGGWRXRRBRRXRRBR
Calculated mass: 9086.0 Da
Observed mass: 9085.8 Da

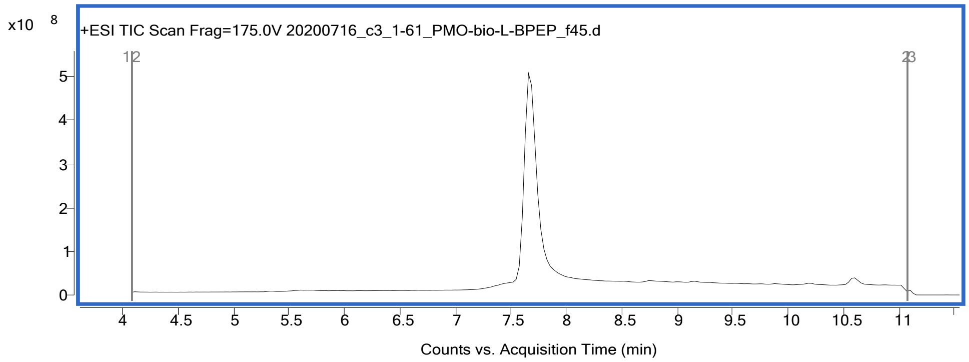

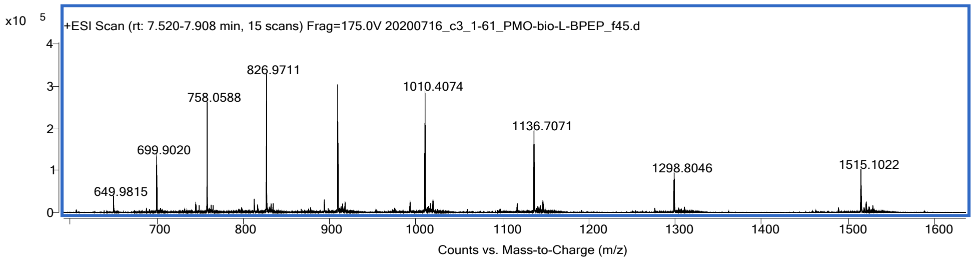

Peptide: PMO-Bio-D-R8
Sequence: Biotin-Lys(PMO)-GGKGGWRRRRRRRR
Calculated mass: 8717.5 Da
Observed mass: 8717.7 Da

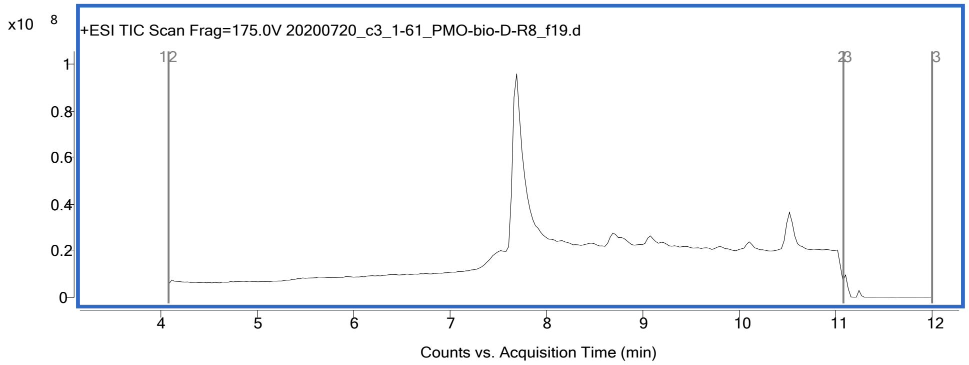

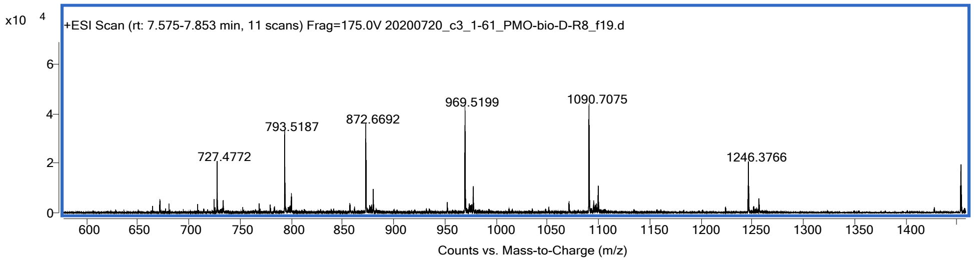

Peptide: PMO-Bio-L-R8
Sequence: Biotin-Lys(PMO)-GGKGGWRRRRRRRR
Calculated mass: 8717.5 Da
Observed mass: 8717.6 Da

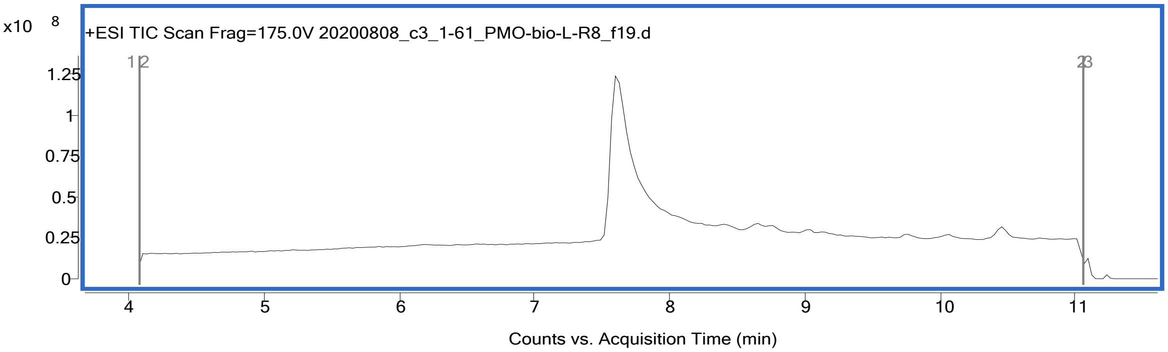

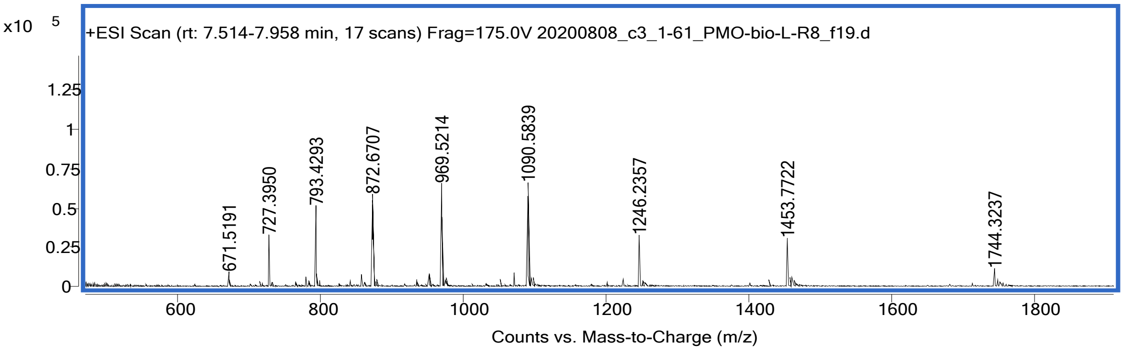
